# WASP and SCAR are evolutionarily conserved in actin-filled pseudopod-based motility

**DOI:** 10.1101/051821

**Authors:** Lillian K. Fritz-Laylin, Samuel J. Lord, R. Dyche Mullins

## Abstract

Diverse eukaryotic cells crawl through complex environments using distinct modes of migration. To understand the underlying mechanisms and their evolutionary relationships, we must define each mode, and identify its phenotypic and molecular markers. Here, we focus on a widely dispersed migration mode characterized by dynamic, actin-filled pseudopods that we call “*α*-motility.” Mining genomic data reveals a clear trend: only organisms with both WASP and SCAR/WAVE—activators of branched actin assembly—make actin-filled pseudopods. While SCAR has been shown to drive pseudopod formation, WASP’s role in this process is controversial. We hypothesize that these genes together represent a genetic signature of *α*-motility, because both are used for pseudopod formation. WASP depletion from human neutrophils confirms that both proteins are involved in explosive actin polymerization, pseudopod formation, and cell migration, and colocalize to dynamic signaling structures. Moreover, retention of WASP together with SCAR correctly predicts *α*-motility in disease-causing chytrid fungi, which we show crawl at >30 μm/min with actin-filled pseudopods. By focusing on one migration mode in many eukaryotes, we identify a genetic marker of pseudopod formation, the morphological feature of *α*-motility, providing evidence for a widely distributed mode of cell crawling with a single evolutionary origin.

## Introduction

Eukaryotic cells move using several distinct modes of locomotion, including crawling and flagella-driven swimming. The stereotyped architecture of flagella and the conservation of their protein components make the evolutionary conservation of cell swimming clear. In contrast, “crawling motility” is a collection of distinct processes whose evolutionary relationships are not well understood (Paluch and Raz, 2013; Lämmermann and Sixt, 2009; Rodriguez et al., 2005). Some crawling cells require dedicated adhesion molecules to make specific, high-affinity contacts with their surroundings, while other cells rely on weaker, nonspecific interactions. Crawling cells also employ different mechanisms to advance their leading edge, either assembling polymerized actin networks to push the plasma membrane forward, or detaching the membrane from the underlying cytoskeleton to form a rapidly expanding bleb. Furthermore, some cell types have been shown to use contractile forces to generate forward movement (Lämmermann et al., 2008; Bergert et al., 2012; Liu et al., 2015). Different cells can also employ different sets of molecules to drive similar modes of crawling. In an extreme example, nematode sperm have evolved a method of crawling in which polymer assembly advances the leading-edge membrane but, in these cells, the force-generating polymer networks are composed of “major sperm protein” rather than actin. Given this variety of crawling behaviors, it is clear that one cannot simply assume that the underlying molecular mechanisms are the same.

The best understood mode of crawling is the slow (1-10 μm/hour) creeping of adherent animal cells, including fibroblasts and epithelial cells (Petrie and Yamada, 2015). These cells move by extending across a surface a sheet-like protrusion called a lamellipodium while gripping substrate molecules using integrins, often clustered into large focal adhesions. Although clinically and physiologically important, this form of adhesion-based crawling is unique to the animal lineage, and largely restricted to molecular “highways” formed by the extracellular matrix.

In contrast, many motile cells—including free-living amoebae and human immune cells—make three-dimensional, actin-filled pseudopods and navigate complex environments at speeds exceeding 20 μm/min (100-1000× faster than fibroblasts) without forming specific molecular adhesions (Buenemann et al., 2010; Butler et al., 2010). Although this mode of fast cell crawling has been called “amoeboid motility,” this term is also used to describe a range of behaviors, including cell motility that relies on membrane blebs rather than actin-filled pseudopods (Lämmermann and Sixt, 2009).

To narrow our focus, we use the term “*α*-motility” specifically to describe cell crawling that is characterized by: (i) highly dynamic three-dimensional pseudopods at the leading edge that are filled with branched-actin networks assembled by the Arp2/3 complex; (ii) fast migration, typically on the order of tens of μm/min; and (iii) the absence of specific, high-affinity adhesions to the extracellular environment. This independence from specific molecular adhesions separates *α*-motility from the adhesion-based motility of fibroblasts and epithelial cells. Furthermore, the use of pseudopods discriminates it from the fast bleb-based motility adopted by fibroblasts in environments that preclude adhesion formation (Liu et al., 2015; Ruprecht et al., 2015). Some organisms using *α*-motility may also employ additional methods of generating forward movement, such as contractility, retrograde flow, and/or blebbing (Lämmermann et al., 2008; Bergert et al., 2012; Yoshida and Soldati, 2006), but here we focus on a single phenotype readily observable in diverse species, including non-model organisms.

Organisms with cells capable of *α*-motility appear throughout the eukaryotic tree, and we hypothesize that this form of locomotion reflects a single, discrete process that arose early in eukaryotic evolution and has been conserved. If this hypothesis is correct, then elements of this ancient process—specific molecules and mechanisms—should be conserved and still associated with cell crawling in distantly related organisms that employ *α*-motility. Such molecular remnants would help to unravel the evolutionary history of cell locomotion, and might enable us to predict the existence of specific modes of motility in poorly characterized species. Identifying genes associated with a process such as *α*-motility is not trivial because the core machinery driving pseudopod formation (e.g. actin and the Arp2/3 complex) is shared with other cellular processes, including some types of endocytosis (Winter et al., 1997). The participation of these proteins in multiple essential processes likely explains their ubiquity in the eukaryotic family tree. Actin, for example, is found in the genomes of all eukaryotes, and based on phylogentic analysis is widely accepted to have been present in the eukaryotic ancestor (Goodson and Hawse, 2002). The Arp2/3 complex is also highly conserved (Beltzner and Pollard, 2004); it is present in the genomes of all sequenced eukaryotes except the parasitic protist *Giardia*, whose lineage either split off before the evolution of the Arp2/3 complex, or lost it, depending on the placement of the root of the eukaryotic tree (Paredez et al., 2011). The ubiquity and multifunctionality of actin and the Arp2/3 complex make them difficult to use as markers for tracing the evolutionary history of *α*-motility.

We therefore turned our attention to upstream regulators of actin assembly. There are a number of “nucleation promoting factors” that stimulate branched actin network assembly by the Arp2/3 complex in response to various upstream cellular signals (Rottner et al., 2010). Some of these Arp2/3 activators are restricted to specific eukaryotic lineages, particularly multicellular animals that evolved the JMY and WHAMM families of Arp2/3 activators, while other activators are more widely distributed (Veltman and Insall, 2010b; Kollmar et al., 2012). For example, WASP and SCAR (also known as WAVE) are widely conserved Arp2/3 activators that respond to different signaling cascades (Rohatgi et al., 1999; Koronakis et al., 2011; Moreau et al., 2000) and promote different levels of Arp2/3 activity (Zalevsky et al., 2001). Multiple published phylogenetic analyses of WASP and SCAR gene families suggest that both genes are ancient and likely to have been present in the eukaryotic ancestor (Veltman and Insall, 2010a; Kollmar et al., 2012). However, neither family is found in all eukaryotic lineages making them appealing candidates for genetic markers of *α*-motility.

SCAR is generally accepted to play a major role in the formation of protrusions used for cell motility (Steffen et al., 2004; Miki et al., 1998; Veltman et al., 2012; Weiner et al., 2006). In contrast, the involvement of WASP genes—particularly the two mammalian homologs WASP and N-WASP—in cell crawling is less clear. N-WASP is ubiquitously expressed in mammals, and is dispensable for lamellipodia or filopodia formation by adherent fibroblasts (Snapper et al., 2001; Lommel et al., 2001; Sarmiento et al., 2008), which has lead many researchers to discount a role for any WASP protein in protrusions or motility (Small and Rottner, 2010; Veltman and Insall, 2010b). Mammalian WASP, on the other hand, is expressed only in blood cells, where it has been shown to be involved in migration and pseudopod formation (Jones et al., 2002; Burns et al., 2001; Badolato et al., 1998; Jones et al., 2013; Shi et al., 2009; Ishihara et al., 2012). Further evidence for a role of WASP in cell migration comes from the handful of papers studying WASP in non-mammalian cells (Veltman et al., 2012; Zhu et al., 2016). (See Table S1 for an annotated bibliography of 29 papers on WASP and N-WASP relating to cell migration, summarized above.) The use of WASP by highly motile cells, but not adherent fibroblasts, may therefore reflect unique requirements for *α*-motility.

To understand the regulation of the actin cytoskeleton during pseudopod formation, we exploit the diversity of organisms that use *α*-motility: by comparing the genomes of many eukaryotes, we find that organisms with genes encoding *both* WASP and SCAR make pseudopods, and organisms that do not build pseudopods have lost either or both Arp2/3 activator. We validate this molecular signature using a negative test (depleting the protein disrupts pseudopod formation in well-studied cells), as well as a positive test (a new prediction of *α*-motility in a little-studied organism). Differentiating *α*-motility from slow/adhesive cell migration helps clarify the confusion over WASP’s importance in cell motility, and shifts the major question from whether WASP *or* SCAR is required for motility in a given single cell type, to how WASP and SCAR work *together* to construct and maintain pseudopods in many species. The retention of WASP and SCAR by organisms that form pseudopods represents the first molecular support, to our knowledge, for a single origin of this widespread form of cell motility in an ancestor of extant eukaryotes (see summary Figure 8).

## Results

### Evolutionary retention of both WASP and SCAR correlates with pseudopod formation

To trace the evolutionary history of *α*-motility, we first determined which sequenced eukaryotic organisms might employ *α*-motility. The obvious structural feature associated with *α*-motility are dynamic, actin-filled pseudopods. In addition to *α*-motility, some organisms use these structures for feeding. Pseudopods used for feeding and α-motility share so many structural and signaling components that, unless specific receptors and/or prey are known to be involved, they are largely indistinguishable (Heinrich and Lee, 2011). Therefore, we combed the literature for references to organisms with cells that form pseudopods for feeding and/or *α*-motility. Eukaryotic phyla fall into at least six large clades, and species with sequenced genomes and that form pseudopods can be found in most (Table 1 and summary Figure 8).

**Table 1.** WAVE and WASP orthologs for indicated species (see (Kollmar et al., 2012) for protein sequences, except for orthologs in *S. rosea*, which we identified via BLAST, with indicated NCBI identifiers. Example references for observations of pseudopods are given. Cases where genes or reference to cells with active pseudopod were not found are indicated by “n.f.” (See also Figure 8.)

| Species | Group | WAVE | WASP | Pseudopod reference |
| --- | --- | --- | --- | --- |
| *H. sapiens* | Animals<br>(opisthokont) | WAVE1, WAVE2,<br>WAVE3 | WASP, N-WASP | (Ramsey, 1972) |
| *M. brevicollis*<br>(Orthologs in *S. rosea*) | Choano-<br>flagellate<br>(opisthokont) | MbWAVE<br>(*S. rosea*:<br>XP_004994362) | MbWASP<br>(*S. rosea*:<br>XP_004997564) | Pseudopod-based feeding has been<br>observed in all choanoflagellate families<br>(Pettitt et al., 2002)<br>(For detailed analysis of pseudopods of<br>*S. rosea* see (Dayel and King, 2014) |
| *C. owczarzaki* | Opisthokont | CoWAVE | CoWASP | (Hertel et al., 2002) |
| *A. nidulans*<br>(Representative<br>dikaryon) | Fungi<br>(opisthokont) | n.f. | EnWASP | n.f. |
| *B. dendrobatidis* | Fungi<br>(opisthokont) | Bad_bWAVE | Bad_bWASP | see Figure 2 |
| *A. macrogynus* | Fungi<br>(opisthokont) | AlmWAVE | AlmWASP_Aalpha,<br>AlmWASP_Abeta,<br>AlmWASP_Balapha,<br>AlmWASP_Bbeta | n.f. |
| *D. discoideum* | Amoebozoan | SCAR1 | DdWASP | (Raper, 1935) |
| *E. histolytica* | Amoebozoan | n.f. | n.f. | n.f. |
| *A. castellanii* | Amoebozoan | AcWAVE | AcWASP_A,<br>AcWASP_B,<br>AcWASP_C | (Bowers and Korn, 1968) |
| *T. trahens*++ | Apusozoa | TctWAVE | TctWASP | (Cavalier-Smith and Chao, 2010) |
| *A. thaliana* | Plant | SCAR1, SCAR2,<br>SCAR3, SCAR4,<br>SCAR-like domain-<br>containing protein | n.f. | n.f. |
| *P. ramorum* | Chromalveolate | n.f. | n.f. | n.f. |
| *B. natans* | Chromalveolate | n.f. | BinWASP_A,<br>BinWASP_B | n.f. |
| *P. falicparum* | Chromalveolate | n.f. | n.f. | n.f. |
| *T. thermophila* | Chromalveolate | n.f. | TtWASP_A,<br>TtWASP_B | n.f. |
| *E. huxleyi* | Rhizarian | n.f. | EmhWasp | n.f. |
| *N. gruberi* | Heterolobosean | NgWAVE_A,<br>NgWAVE_B | NgWASP_A,<br>NgWASP_B,<br>NgWASP_C,<br>NgWASP_D | (Fulton, 1970) |
| *G. lamblia* | Diplomonad | n.f. | n.f. | n.f. |
| *T. vaginalis* | Parabasalid | Tv_aWAVE_A,<br>Tv_aWAVE_B,<br>Tv_aWAVE_C,<br>Tv_aWAVE_D,<br>Tv_aWAVE_E,<br>Tv_aWAVE_F,<br>Tv_aWAVE_G,<br>Tv_aWAVE_H,<br>Tv_aWAVE_I,<br>Tv_aWAVE_J,<br>Tv_aWAVE_K | Tv_aWasp_A,<br>Tv_aWasp_B | (Kusdian et al., 2013) |

On to this map of the phylogenetic distribution of pseudopods, we overlaid the conservation of WASP and SCAR/WAVE genes, using a recently published, manually curated database of nucleation promoting factors from genomes spanning eukaryotic diversity (Kollmar et al., 2012). Multiple analyses have concluded that both WASP and SCAR were present in the last common ancestor of eukaryotes (Veltman and Insall, 2010a; Kollmar et al., 2012), and therefore argue that a lack of either gene reflects loss during evolution.

To understand whether these gene loss events reveal a significant pattern, we compared the conservation of individual nucleation promoting factors across large evolutionary distances with the ability to assemble pseudopods. We identified a correlation between the conservation of WASP and SCAR and pseudopod formation (Table 1 and summary Figure 8).

For example, *no* plant cells build pseudopods and *no* sequenced plant genomes contain a WASP ortholog. Similarly, multicellular fungi—the dikarya—lack SCAR and are also not known to build pseudopods. Conversely, almost all sequenced genomes of Amoebozoan species (including dictyostelids) encode orthologs of WASP and SCAR, and almost all move with the help dynamic, actin-rich pseudopods. A potential counterexample is the amoeba *Entamoeba histolytica* that lacks *both* WASP and SCAR, yet forms Arp2/3-dependent phagocytic “food cups” to engulf bacteria (Babuta et al., 2015). The absence of both genes indicates that *Entamoeba* must use another Arp2/3 activation system for protrusions, an idea supported by its use of blebs and not pseudopods for motility (Maugis et al., 2010). But the most glaring exception to the correlation was a pair of little-studied species of chytrid fungi that retain both nucleation promoting factors, but are not known to build pseudopods: *Allomyces macrogynus* and *Batrachochytrium dendrobatidis*.

We took a two-pronged approach to testing our hypothesis that retention of WASP together with SCAR serves as a molecular signature of pseudopod formation. First, we took the more traditional approach and confirmed that both genes are involved in pseudopod formation in mammalian cells. We followed this with an evolution-based approach by verifying the ability of this molecular signature to predict the capacity for pseudopod formation in chytrid fungi.

### WASP and SCAR localize to the same dynamic arcs within pseudopods of human neutrophils

Our evolutionary evidence indicates that WASP and SCAR may *both* be required to build pseudopods. To test this hypothesis directly, we turned to human cell lines capable of forming pseudopods. HL-60 cells are derived from an acute myeloid leukemia (Collins et al., 1977) and retain many features of hematopoietic cells, including expression of hematopoietic WASP and the capacity to differentiate into fast-migrating neutrophils with dynamic pseudopods (Collins et al., 1978).

To follow the dynamics of WASP localization in live cells, we created an HL-60 line stably expressing full-length WASP fused at the N-terminus to the red florescent protein TagRFP-T. By confocal fluorescence microscopy, TagRFP-WASP concentrates in two distinct locations within migrating HL-60 cells: punctate foci distributed throughout the cell and a broad zone near the leading edge (Figure S1A).

Others have shown that the SCAR regulatory complex localizes to fast-moving, anterograde **“**waves” that break against the leading edge of actively migrating HL-60 cells (Weiner et al., 2007). This localization pattern is most easily observed using total internal reflection fluorescence (TIRF) microscopy, which illuminates a ∼100 nm thick region of the cell near the ventral surface (Axelrod, 1981). Using TIRF microscopy on rapidly migrating HL-60 cells, we observed that TagRFP-WASP concentrates near the leading edge in linear arcs that move in an anterograde direction, similar to previously observed patterns of the SCAR regulatory complex (Weiner et al., 2007).

To see whether WASP and the SCAR travel together in the same waves, we introduced TagRFP-WASP into cells expressing YFP-Hem1, a core component of the SCAR regulatory complex (Weiner et al., 2007). TIRF microscopy of these cells revealed that WASP and the regulatory SCAR complex move together in the same dynamic, linear arcs (Figure 1A-B, Figure S1B, and **Video 1**). Interestingly, however, the localization patterns of the two are not identical, an observation confirmed by quantifying WASP and SCAR localization across the leading edge (Figure S1C). Spinning disk confocal microscopy indicates that WASP and SCAR colocalize throughout the growing pseudopods, not only at the ventral surface (Figure 1C). Within the resolution limits of our imaging, the localization patterns move together, with neither protein consistently leading the other (Figure S1B and **Video 1**). This dynamic localization pattern suggests that both WASP and SCAR activate the Arp2/3 complex in leading-edge pseudopods, promoting assembly of the branched actin networks required for membrane protrusion.

**Figure 1:**
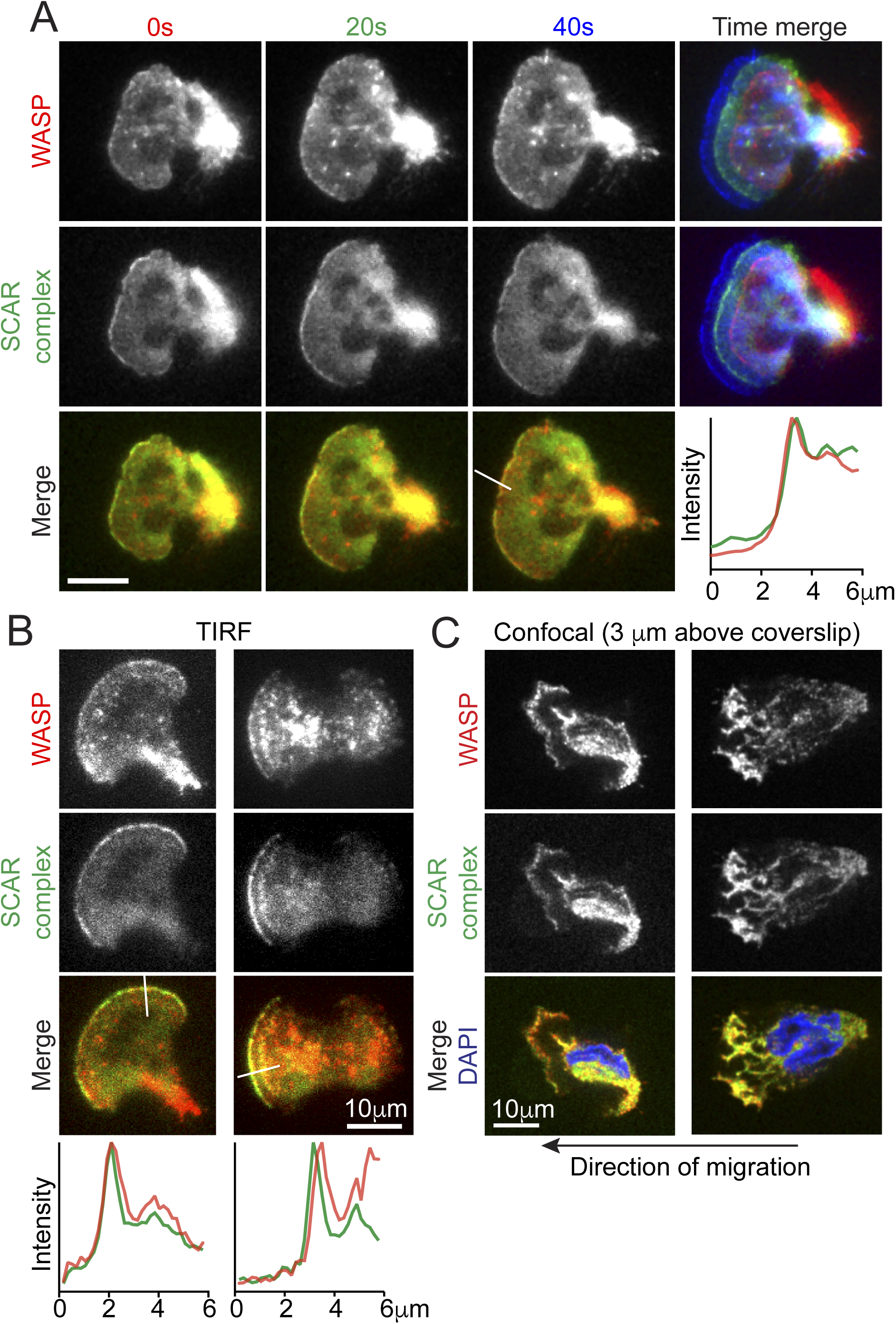
WASP colocalizes with the SCAR complex at the leading edge of neutrophils. Microscopy of HL-60 cells expressing TagRFP-WASP and Hem-1-YFP, a component of the SCAR regulatory complex. For all cells, the direction of migration is to the left. **(A)** TIRF images of live HL-60 cells. Top: WASP localization in three sequential time points, overlaid on the far right (0 seconds in red, 20 seconds in green, and 40 seconds in blue). Middle: same sequence of images, but for Hem-1. Bottom: overlay of WASP and Hem-1 at each time point. The plot shows line scans of normalized fluorescence intensity of WASP (red) and Hem-1 (green). The location for generating the line scans is shown in the adjacent image. **(B)** Two additional examples of live HL-60 cells in TIRF and corresponding line scans. **(C)** Spinning disk confocal images of two fixed HL-60 cells, showing an axial slice through the middle of thick pseudopods. The slices shown were taken 3 μm above the coverslip. See also Figure S1 for additional images and kymographs, and **Video 1**.

### WASP participates in pseudopod assembly in neutrophils

To investigate whether WASP is involved in pseudopod assembly by HL-60 cells, we generated anti-WASP small hairpin RNAs (shRNAs), expression of which resulted in a >90% reduction of WASP protein, but no obvious change in WAVE2 (Figure 2A). We next examined whether WASP-depleted (WASP-KD) cells can form pseudopods. In a gradient of chemoattractant (the peptide fMet-Leu-Phe), wildtype HL-60 cells become strongly polarized with broad, actin-rich pseudopods used to rapidly move toward the source of chemoattractant (Figure 2B, and **Video 2**). Compared to control, 50% fewer WASP-KD cells formed pseudopods (Figure 2B-C). Despite numerous attempts, we never succeeded in developing WASP-KD cell lines in which this phenotype was 100% penetrant. Although it is possible that this is due to residual WASP protein, this seems an insufficient explanation because the WASP-KD cells still capable of forming pseudopods are also aberrant (see below). Moreover, WASP knockout mice also show only a partial defect in the gross cell motility *in vivo* (Snapper et al., 2005).

**Figure 2.**
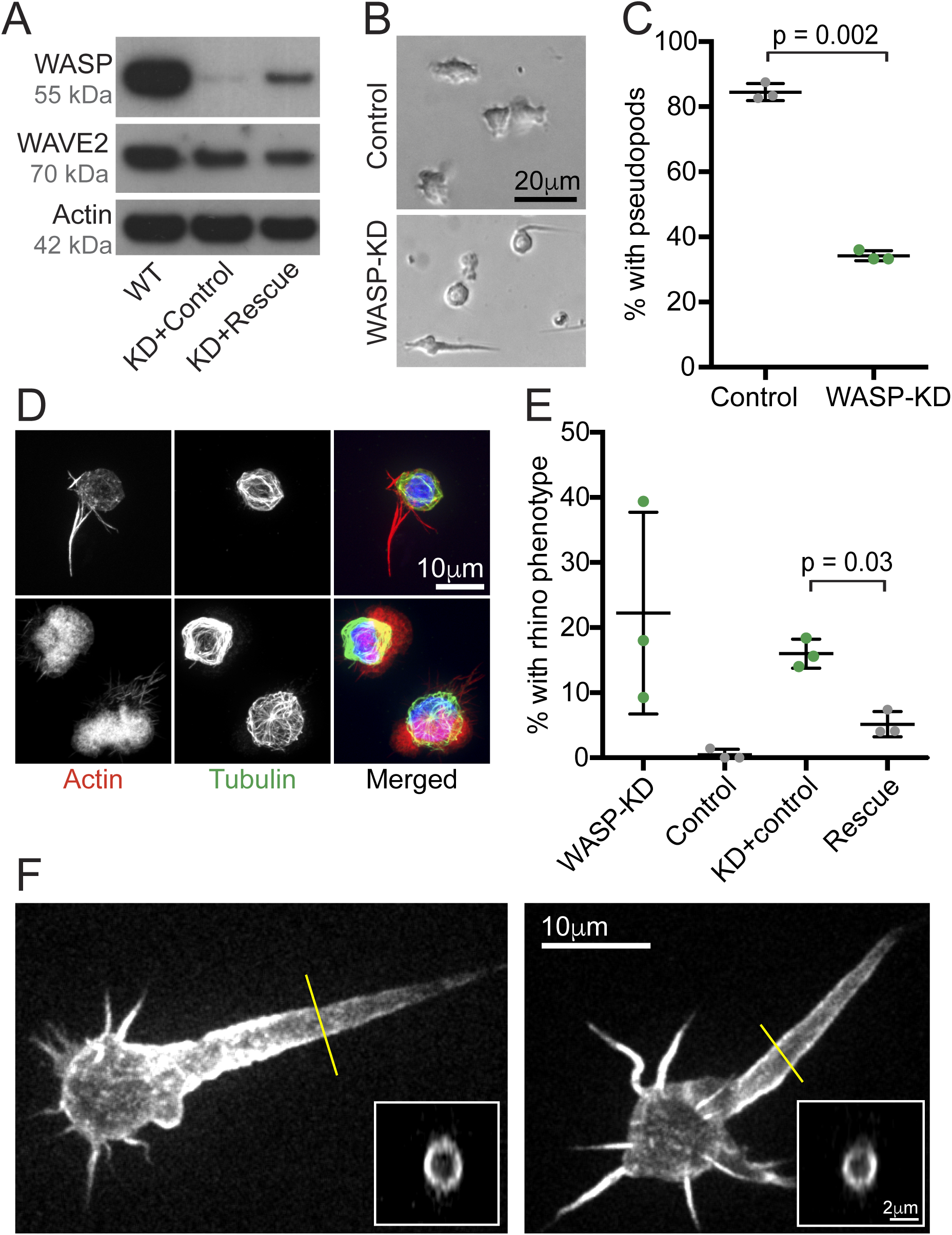
WASP is crucial for pseudopod formation of neutrophils. **(A)** Western blots showing WASP and WAVE2 expression in control HL-60 cells, cells expressing shRNA to WASP and exogenous WASP with three silent mutations in the region corresponding to the shRNA (KD+rescue), or cells expressing anti-WASP shRNA and empty vector rescue control (KD+control). Approximately equal amounts of total protein were loaded in each lane, confirmed using actin as a loading control. **(B)** Brightfield images of control and WASP knockdown cells. **(C)** Knockdown of WASP using shRNA reduces the percentage of cells that have pseudopods. Means and standard deviations (bars) of three biological replicates (dots) with >135 cells total; p value from two-tailed paired t-test. **(D)** Immunofluorescence of control and WASP-KD HL-60 cells showing microtubules (green, antibody-stained), actin filaments (red, phalloidin-stained), and DNA (blue, DAPI). Note the signature rhino phenotype in the WASP-KD cell. See also Figure S2. **(E)** Rescue of rhino protrusion phenotype by expression of shRNA insensitive WASP as described in (A). Note: no wildtype cells were observed to exhibit the rhino phenotype. Means and standards deviation (bars) of three biological replicates (dots) with >130 cells total; p value from two-tailed paired t-test. **(F)** Maximum projections of spinning disk confocal stacks of living HL-60 cells with fluorescent probe specific for polymerized actin (mCherry fused to calponin homology domain of Utrophin, Utr261). Insets are cross sections through the rhino horns at positions indicated by yellow lines, confirming that they are hollow with a shell of actin. See also Figure S2 for time lapse of right hand cell showing the dynamics of the rhino horn protrusion.

In addition to the defect in pseudopod formation, approximately 20% of WASP-KD cells formed large protrusions that taper to a point, reminiscent of a rhinoceros horn (Figure 2B,D,F Figure S2, and **Video 2** WASP-KD cells 11, 32, 33, 39 and 42, for example). To verify its specificity, we rescued this “rhino” phenotype by expressing a functional WASP containing three silent mutations in the sequence targeted by to the shRNA (Figure 2E). Additionally, a second shRNA that targets a separate region of the WASP gene resulted in a significantly smaller effect on both WASP expression and the number of cells with the rhino phenotype (not shown). Immunofluorescence combined with phalloidin staining of polymerized actin revealed that the aberrant rhino protrusions contain actin filaments but lack microtubules (Figure 2D). The expression of a probe specific for polymerized actin (mCherry fused to calponin homology domain of Utrophin, Utr261 (Burkel et al., 2007)) revealed a highly dynamic and, surprisingly, hollow actin filament network inside the protrusions (Figures 2F and S2). This distribution—enriched near the membrane but depleted from the core of the protrusion—is more reminiscent of cortical actin networks than of filopodia, which are packed tight with actin bundles (Tilney et al., 1973).

Because some WASP family proteins contribute to endocytosis (Benesch et al., 2005; Merrifield et al., 2004; Naqvi et al., 1998), we investigated whether the defects in WASP-KD cells are caused by reduction of endocytosis. In undifferentiated HL-60s, we observed no difference in transferrin receptor endocytosis and recycling between WASP-KD and control cells (Figure S3E). After differentiation into cells capable of making pseudopods, WASP-KD HL-60s actually showed increased surface receptor densities (Figure S3D), receptor internalization (Figure S3B), as well as receptor recycling (Figure S3C) compared to control cells. Therefore, we cannot attribute the WASP-KD phenotypes simply to a curtailment of endocytosis activity.

### WASP-depleted neutrophils polymerize less actin in response to chemoattractant

Addition of chemoattractant to non-polarized (quiescent) HL-60 cells induces a burst of actin polymerization that drives polarization and pseudopod formation, nearly doubling the cell’s polymerized actin content within 30 seconds of stimulation. This response is already known to depend on the activity of the SCAR regulatory complex (Weiner et al., 2006). To determine what role WASP might play in this explosive actin assembly, we synchronized pseudopod formation by stimulating populations of quiescent HL-60s with fMLP, fixed and stained the cells with phalloidin at different time points, and analyzed total polymerized actin content in each cell by confocal microscopy and fluorescence-activated cell sorting (FACS) (Figure 3A–B). In the absence of chemoattractant, the amount of polymerized actin in quiescent WASP-KD cells was roughly equal to that in control cells. However, as reported for SCAR-depleted cells (Weiner et al., 2006), WASP-KD cells had greatly reduced actin polymerization at both short (30 seconds) and long times (3 min) following stimulation. This reduced actin polymerization indicates that WASP, like SCAR, is central to the explosive actin polymerization required for cell polarization and subsequent pseudopod formation.

**Figure 3.**
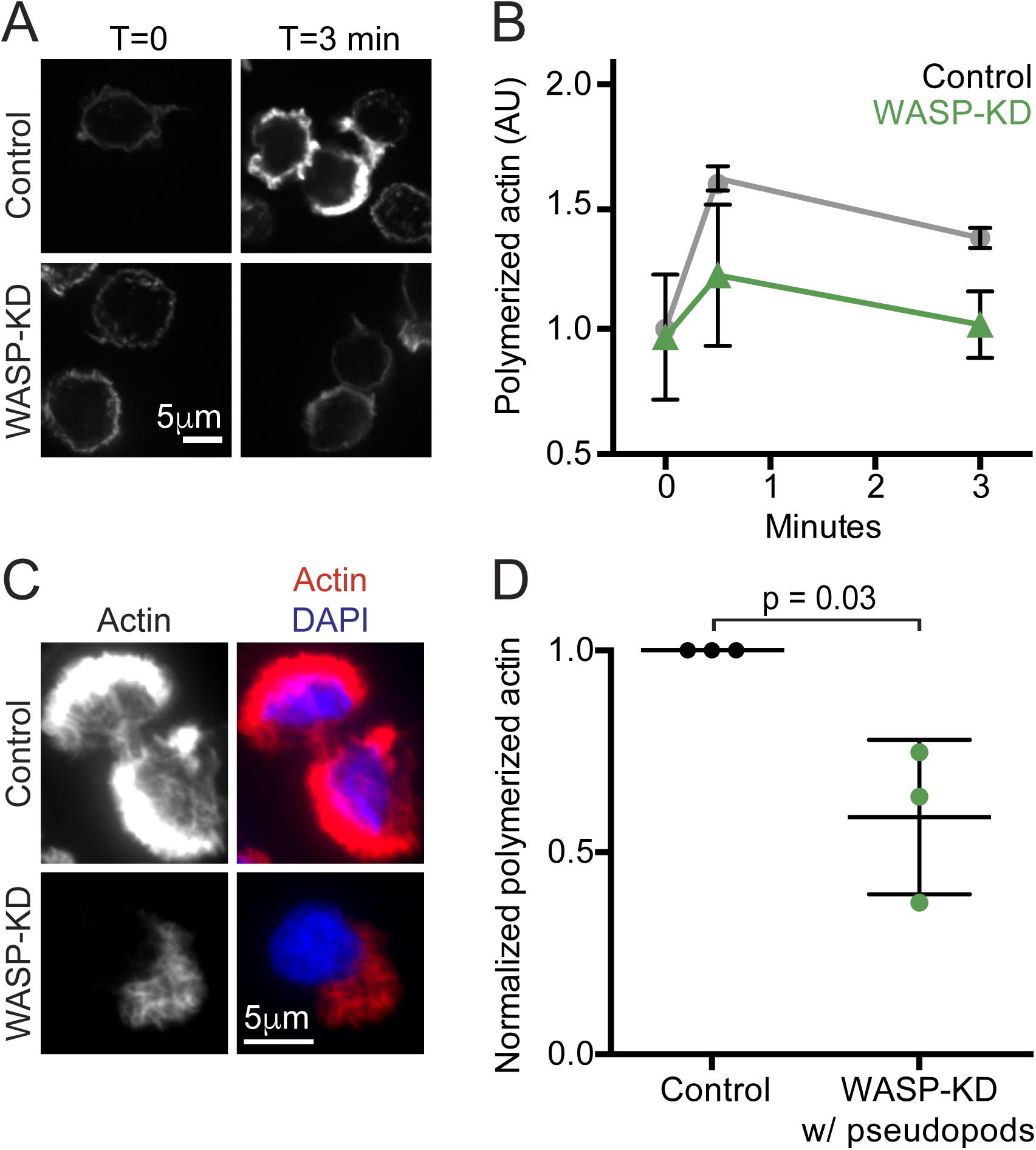
WASP is crucial for explosive actin polymerization during pseudopod formation in neutrophils. **(A)** Spinning disk confocal images of polymerized actin of control and WASP-KD HL-60 cells after stimulation for the indicated time with chemoattractant (20 nM fMLP), stained with florescent phalloidin**. (B)** FACS quantification of actin polymerization in control (grey squares) and WASP-KD HL-60 cells (green triangles) after stimulation for the indicated time with chemoattractant, stained with florescent phalloidin. 10,000 cells were counted for each sample, and averages normalized to time zero for control cells within each experiment to control for detector variability. Symbols and error bars are means and standard deviations from 3 biological replicates. (**C**) Example spinning disk confocal images of pseudopod-forming WASP-KD and control cells, with polymerized actin stained with phalloidin (red) and DNA stained with DAPI (blue). (**D**) Quantitation of phalloidin staining shown in (C). Only cells with pseudopods were analyzed. Mean pixel value for z-projections of image stacks was measured for each cell, then the average background pixel value was subtracted. Means and standard deviation (bars) from 3 biological replicates (dots), with >150 cells total; p value from one-tailed paired t-test.

To understand more about the pseudopods formed by some WASP-KD cells, we analyzed confocal images of hundreds of individual phalloidin-stained cells. Quantitation of the actin content in the subset of WASP-KD cells that make pseudopods compared to control cells revealed that even WASP-KD cells that appear to make “normal” pseudopods contain about half the quantity of polymerized actin (Figure 3C-D).

### WASP depletion impairs neutrophil motility

To determine the effect of WASP depletion on cell locomotion, we imaged HL-60 cells migrating through a chemoattractant gradient in a 5 μm tall glass chamber (Millius and Weiner, 2010). Tracking individual cells revealed a severe migration defect in WASP-KD cells (reported mean ± standard deviation of three biological replicates): while control cells moved at 12 ± 0.8 μm/min, WASP-KD cells averaged 5.5 ± 1.5 μm/min, and cells with rhino protrusions were almost completely immotile moving at 1.7 ± 0.4 μm/min (Figure 4A–B and **Video 2**). This motility defect is not limited to the rhino cells; when these cells are excluded from the analysis, we still observed a significantly reduced speed (6.7 ± 0.8 μm/min) compared to control (Figure 4B). We did not observe an effect of WASP depletion on directional persistence (Figure 4C).

**Figure 4.**
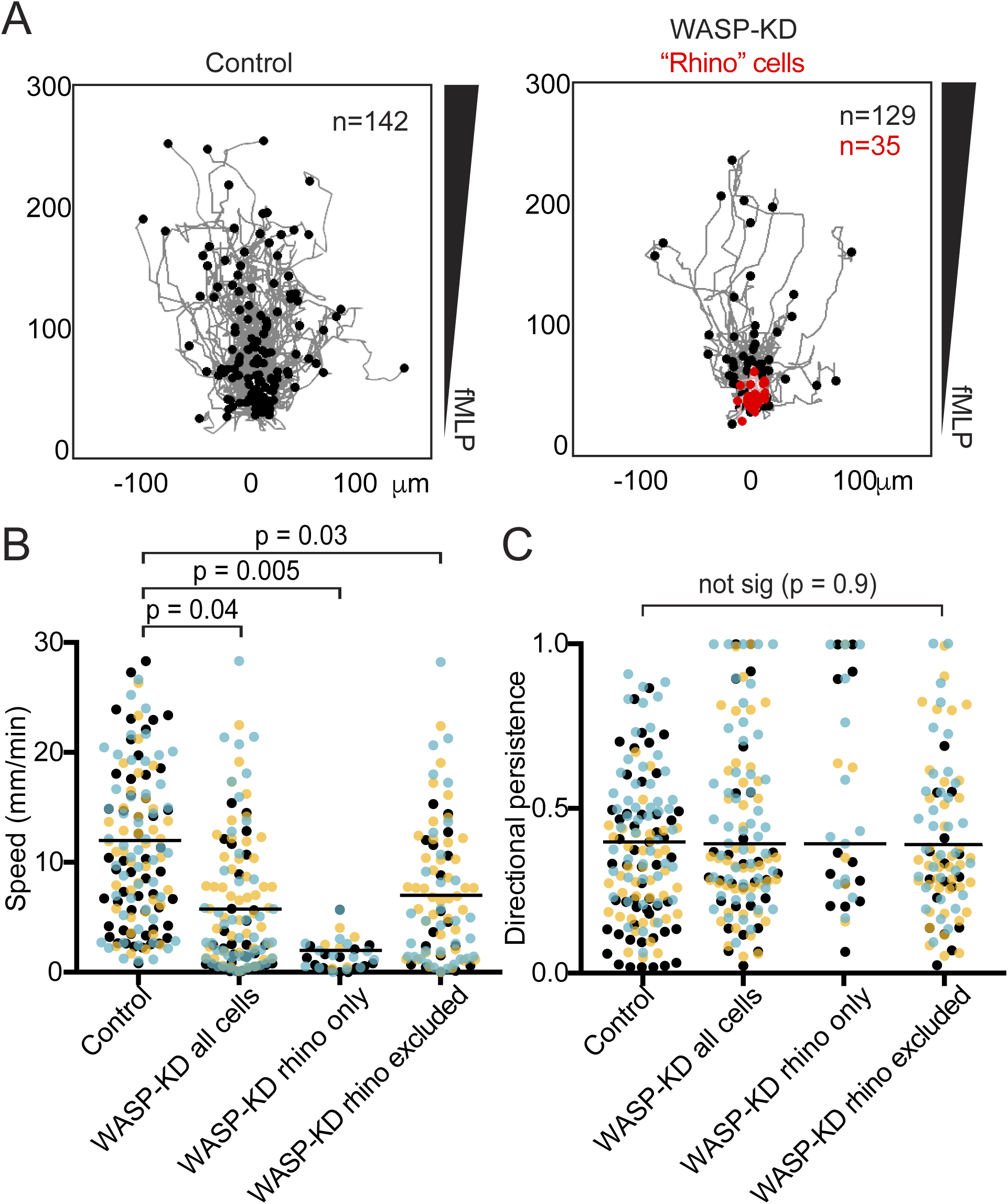
WASP is crucial for neutrophil motility. **(A)** Worm plots showing the tracks of cell migrating up a fMLP chemoattractant gradient: control cells (left), WASP-KD cells (right), with cells that exhibit the rhino phenotype in red. Cells were imaged 20 min, and migration paths overlaid, with time zero at (0,0). The endpoint of each cell’s path is shown with a dot. See also **Video 2. (B)** Depletion of WASP protein leads to reduced cell speed. The average instantaneous speed for each cell in (A) is plotted as a dot, color-coded by biological replicate to highlight the consistency from experiment to experiment. The average of the three replicates is displayed as a horizontal line; p values from two-tailed paired t-test. (**C**) Reduction of WASP protein leads to no significant change in directional persistence (the ratio of the Euclidean distance to the accumulated distance) of cells tracked in (A).

Because the chamber we used to measure directional migration is not pre-coated with fibronectin (or any other specific molecule), we doubt this migration defect is due to an integrin-mediated adhesion defect. We confirmed this by directly testing adhesion to fibronectin-coated surfaces found no significant difference between WASP-KD and control cells (Figure S3A). We conclude that HL-60 cells use WASP, along with SCAR (Weiner et al., 2006), for normal pseudopod formation and efficient *α*-motility.

### WASP and SCAR genes predict pseudopod formation by chytrid fungi

A potential exception to the tight correlation between actin-rich pseudopods and the genomic retention of WASP and SCAR were two deeply branching, and little-studied, species of fungi; the chytrids *Allomyces macrogynus* and *Batrachochytrium dendrobatidis* (*Bd*). These chytrid species contain genes encoding both WASP and SCAR, but have not been reported in the literature to migrate using pseudopods. We were, however, able to find references to pseudopod formation by unsequenced infectious species related to *A. macrogynus* (*Catenaria anguillulae*), which may employ these structures for motility across the surface of its target host (Deacon and Saxena, 1997; Gleason and Lilje, 2009). However, because chytrid fungi are not a monophyletic group, but rather comprise multiple deeply branching clades that are estimated to have diverged around 800 million years ago (Stajich et al., 2009) (James et al., 2006), one cannot assume that distantly related species share this capacity. Therefore, we used Bd as a predictive test of our hypothesis that WASP and SCAR genes represent a marker for *α*-motility.

Like other species of chytrid fungi, the lifecycle of Bd has two stages: a large (10–40 μm) reproductive zoosporangium, which releases a host of small (3–5 μm), motile, flagellated zoospore cells (Longcore et al., 1999; Berger et al., 2005). These infectious zoospores can form cysts beneath the skin of an amphibian host that develop into new zoosporangia to complete the life cycle (Berger et al., 2005). We searched for *α*-motility in Bd zoospores because, unlike the sessile cyst and zoosporangium, these free-swimming flagellates lack a cell wall and have been reported to assume non-uniform shapes with dense “cytoplasmic extensions” (Longcore et al., 1999).

To restrict the fast-swimming Bd zoospores to the imaging plane, we adhered zoospores to Concalavin-A coated glass. In initial experiments, we observed only a small fraction (<1%) of zoospores forming pseudopod-like protrusions. The rarity of pseudopod-forming cells made us suspect that *α*-motility might only occur during a short phase of the life cycle. We therefore enriched for cells of the same age by washing zoosporangia to remove previously released zoospores and collecting flagellates released during the subsequent two hours.

During the first 6 hours after release from the zoosporangium, ~40% of zoospores create dynamic pseudopod-like protrusions (Figure 5A–B, Figure S4A, and **Video 3**) that extend from the cell body at a rate of 25 ± 9 μm/min (Figure 5C), consistent with speeds expected for pseudopods (Chodniewicz and Zhelev, 2003; Zhelev et al., 2004). Unlike blebs, these cellular protrusions are not spherical but irregularly shaped and amorphous—similar to the actin-rich pseudopods of amoebae and neutrophils.

**Figure 5.**
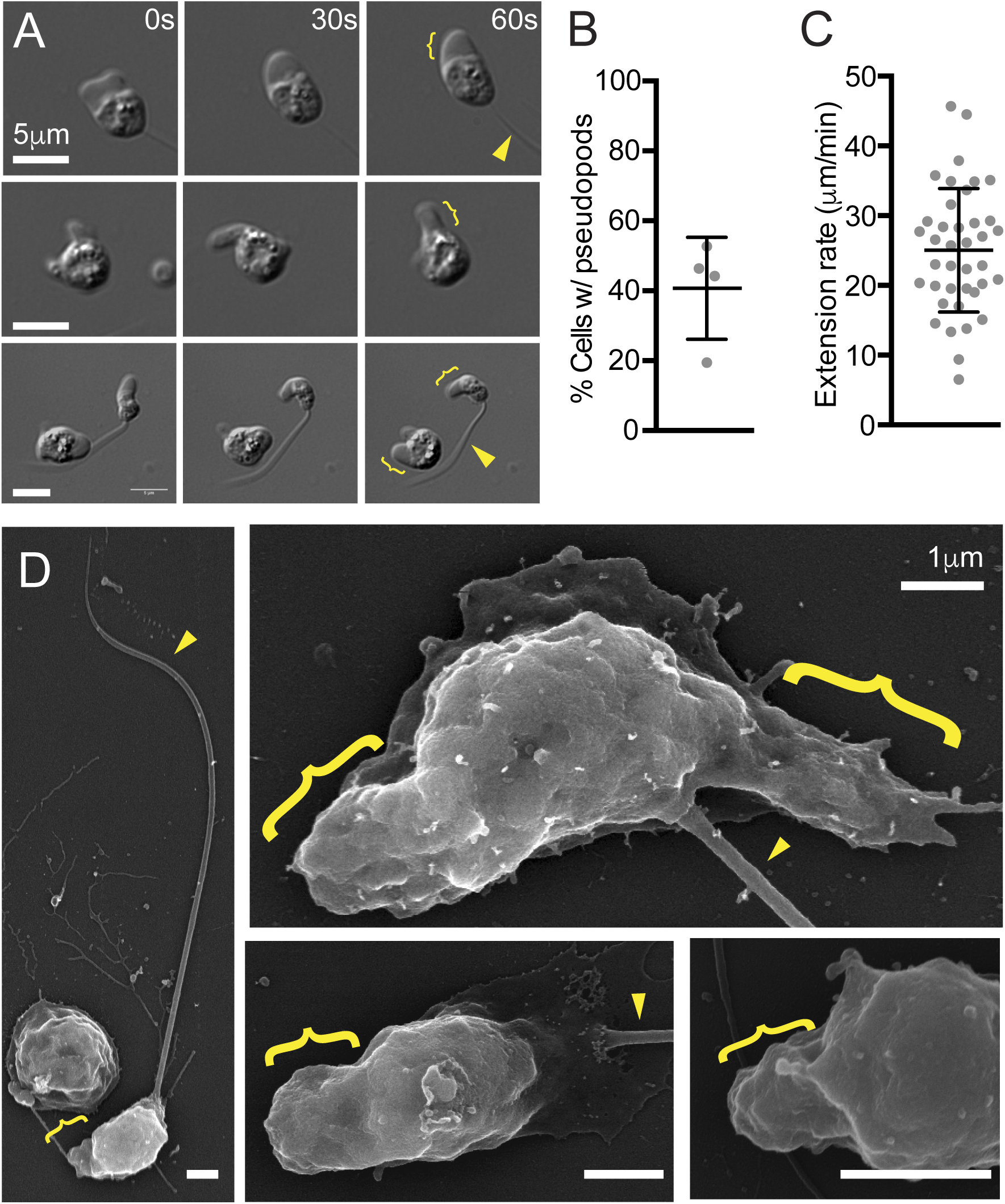
Genomic retention of both WASP and SCAR correctly predicts pseudopod formation by the infectious chytrid fungus *B. dendrobatidis*. Throughout figure, pseudopods are indicated by brackets and flagella by arrowheads. **(A)** Time lapse showing examples of dynamic pseudopods from chytrid cells with (middle) and without a flagellum (middle), or one cell of each (bottom). See also **Video 3**. (**B**) Percentage of cells with pseudopods within first six hours after release from zoosporangia. Mean and standard deviation (bars) of four biological replicates (dots), with 3782 cells total. (**C**) Pseudopod extension rates. Mean and standard deviation (bars) of the individual values (dots) combined from three biological replicates. **(D)** Scanning electron micrographs of fixed chytrid zoospores. See also Figure S4B for more examples.

To ensure that these crawling cells were not contaminating organisms, we obtained independently isolated Bd cultures from three different laboratories, and observed similar pseudopods in each.

To better investigate the morphology of these tiny pseudopods, we performed scanning electron microscopy on fixed cells (Figure 5D and S4B), and observed a similar proportion of flagellated zoospores with one or more thick protrusions. Each protrusion was about 1 μm long and 1 μm wide, and many appeared to be composed of multiple discrete terraces. (Figure 5D and S4B).

### Chytrid pseudopods contain actin and require Arp2/3 activity

Using our assay to image chytrid zoospores, we next investigated whether extension of Bd pseudopods is driven by assembly of branched actin networks, as in other cells crawling using *α*-motility. We first fixed the cells to preserve the actin cytoskeleton and then stained them with fluorescent phalloidin, revealing a thin shell of cortical actin surrounding the cell body and a dense network of filamentous actin filling the pseudopod (Figure 6A).

**Figure 6.**
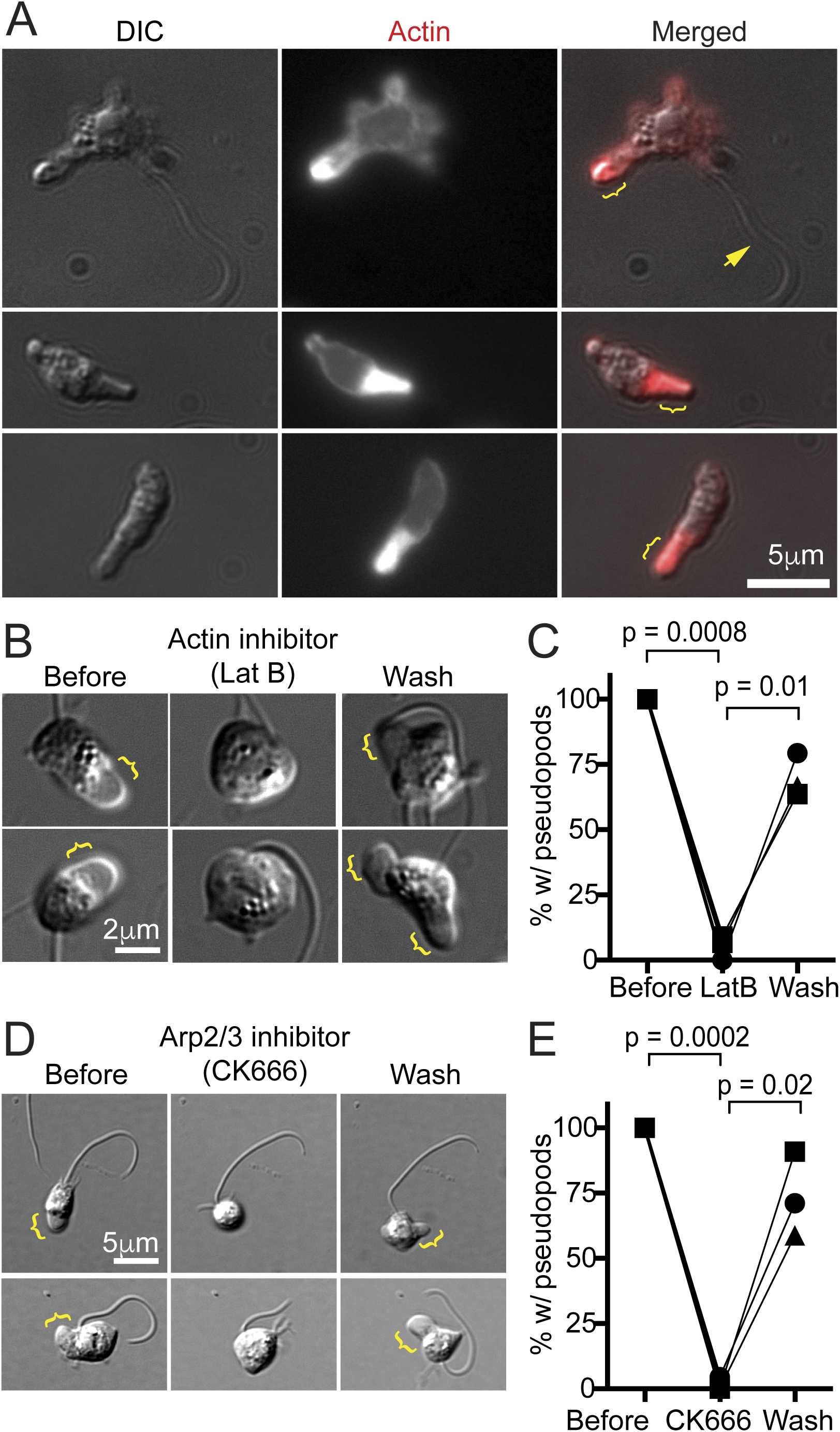
Chytrid pseudopods are actin filled and require both actin polymerization and Arp2/3 activity. **(A)** Fixed chytrid cells with and without a flagellum (arrow). Staining with fluorescent phalloidin reveals a thin shell of cortical actin surrounding the cell body and a dense network of polymerized actin filling the pseudopods (brackets). **(B)** Two examples of chytrid cells with pseudopods that lose them when treated with 10 nm latrunculin B, an inhibitor of actin polymerization. Dynamic pseudopods (brackets) return after drug washout. **(C)** Quantitation of reversible inhibition of pseudopods by latrunculin B. Only cells that were making pseudopods before treatment and that were not washed away during the experiment were counted. Symbols are means from three biological replicates, each with at least 29 cells; p values from two-tailed paired t-test. **(D)** Two examples of chytrid cells with pseudopods that lose them when treated with 10 μm CK-666, an inhibitor of Arp2/3 activity. Pseudopods (brackets) return after drug washout. **(E)** Quantitation of reversible inhibition of pseudopods by CK-666. Only cells that were making pseudopods before treatment and that were not washed away during the experiment were counted. Symbols are means from three biological replicates, each with at least 17 cells; p values from two-tailed paired t-test.

To test whether actin-filled chytrid pseudopods require actin polymerization, we treated zoospores with latrunculin, a small molecule that sequesters actin monomers and inhibits growth of actin polymers. Within minutes of adding 10 nM latrunculin B, nearly all pseudopods ceased growing and/or disappeared (Figure 6B–C). This affect was reversed within an hour after removing the drug.

To determine whether assembly of the pseudopodial actin network requires the nucleation and branching activity of the Arp2/3 complex, we incubated zoospores with CK-666, a small molecule that inhibits actin nucleation by mammalian and fungal Arp2/3 complexes (Nolen et al., 2009). Addition of 10 μM of CK-666 reduced the number of cells with active protrusions by nearly 100%, an effect reversed by washing out the drug (Figure 6D–E). These experiments reveal that protrusion of Bd pseudopods requires Arp2/3-dependent actin assembly.

### Chytrid zoospores use pseudopods for *α*-motility

Although pseudopod-forming Bd cells adhere tightly to glass surfaces coated with Concalavin-A, they were not able to move or swim away from the site of initial attachment, and other coatings did not promote any form of attachment (including collagen, fibronectin, and human keratin, not shown). Several types of animal cells are known to migrate without specific molecular adhesions in confined environments (Lämmermann et al., 2008; Liu et al., 2015; Ruprecht et al., 2015). To test whether Bd zoospores might also be capable of migration in confined environments, we sandwiched cells between two uncoated glass coverslips, held apart by 1 μm diameter glass microspheres, and observed rapidly migrating cells (Figure 7, **Video 4**). Obviously migrating cells had an average instantaneous speed of 19 ± 9 μm/min, with individual cells averaging speeds over 30 μm/min (Figure 7C), consistent with the rates of pseudopod extension described above (Figure 5C). The trajectories of these cells appeared fairly straight (Figure 7B), with an average directional persistence of 0.61 ± 0.25 (Figure 7D).

**Figure 7.**
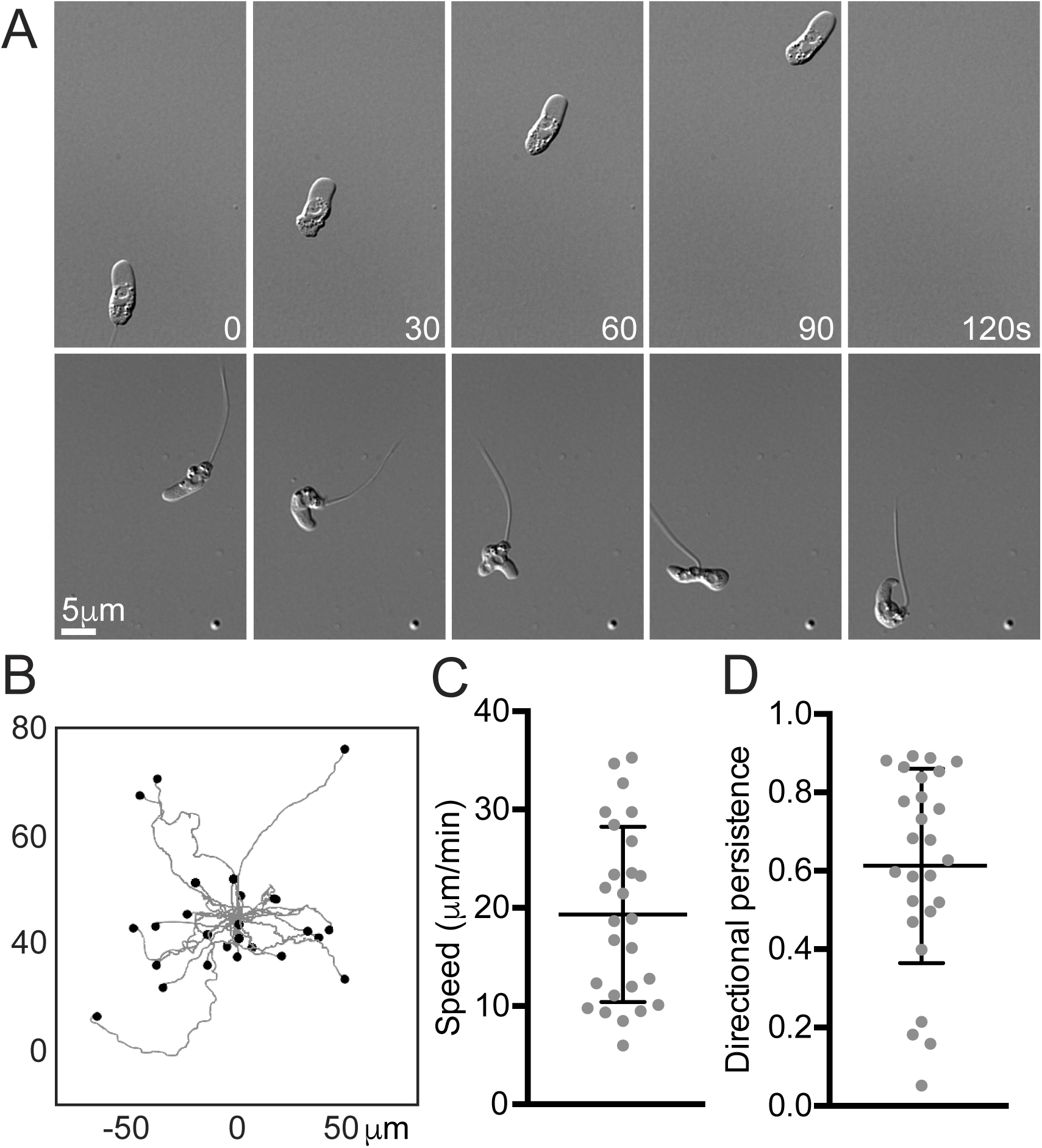
Genomic retention of both WASP and SCAR correctly predicts *α*-motility in the infectious chytrid fungus *B. dendrobatidis*. **(A)** Example chytrid zoospores with and without a flagellum migrating when confined between 1 μm spaced glass coverslips. See also **Video 4**. **(B)** Worm plots showing the tracks of 26 migrating chytrid zoospores, with migration paths overlaid, with time zero at (0,0). The endpoint of each cell’s path is shown with a dot. Only those cells obviously migrating were tracked, and for the duration of their movement. **(C)** Average instantaneous speed of cells tracked in (B). Mean and standard deviation (bars) of the individual values (dots) combined from three biological replicates. **(D)** Directional persistence (the ratio of the Euclidean distance to the accumulated distance) of cells tracked in (B). Mean and standard deviation (bars) of the individual values (dots) combined from three biological replicates.

Some pseudopod-forming zoospores retained flagella, while other cells had clearly lost or resorbed their flagella and strongly resembled free-living amoebae (Figure 7A and **Videos 3**–**4**). We also observed cells switching from crawling to flagellar motility and vice versa, as well as cells rapidly retracting their flagellar axonemes into the cell body (**Video 5**).

## Discussion

Our results reveal that, across eukaryotic phyla, cells that construct actin-rich pseudopods and undergo fast, low-adhesion crawling have retained the Arp2/3 complex as well as two distinct activators of its actin nucleation activity: WASP and SCAR/WAVE. This finding is well supported by a recent paper implicating both WASP and SCAR in *C. elegans* neuroblast cell migration (Zhu et al., 2016). In that system, the phenotype of SCAR mutants is enhanced by loss of WASP, and both WASP and SCAR are found at the leading edge of migrating neuroblasts *in vivo*. Our hypothesis is also consistent with studies of myoblast cell fusion events during muscle formation: like pseudopods, myoblast protrusions are actin-filled force generating machines that require both WASP and SCAR (Sens et al., 2010).

Organisms without the capacity to crawl using pseudopods turn out to have lost one or both of these nucleation-promoting factors (Table 1 and Figure 8). The presence of genes encoding both WASP and SCAR, therefore, provides a molecular correlate for a suite of behaviors that we call “*α*-motility.” The conservation and phylogeny of WASP and SCAR indicate that both were present in a common ancestor of living eukaryotes (Kollmar et al., 2012; Veltman and Insall, 2010a). The power of the co-conservation WASP and SCAR as a genomic marker with the ability to identify cryptic pseudopod-forming organisms, together with cell-biology evidence that both WASP and SCAR are required for *α*-motility in well-studied organisms, argues that this widespread behavior arose from single, ancient origin. It is formally possible that *α*-motility did not have a single evolutionary origin, but that scenario would require both WASP and SCAR to be co-opted *together* for pseudopod assembly multiple times during eukaryotic evolution. Because WASP and SCAR are only two of a large number of Arp2/3 activators (Rottner et al., 2010), we have no reason to believe that motility would repeatedly converge on these two in particular.

**Figure 8.**
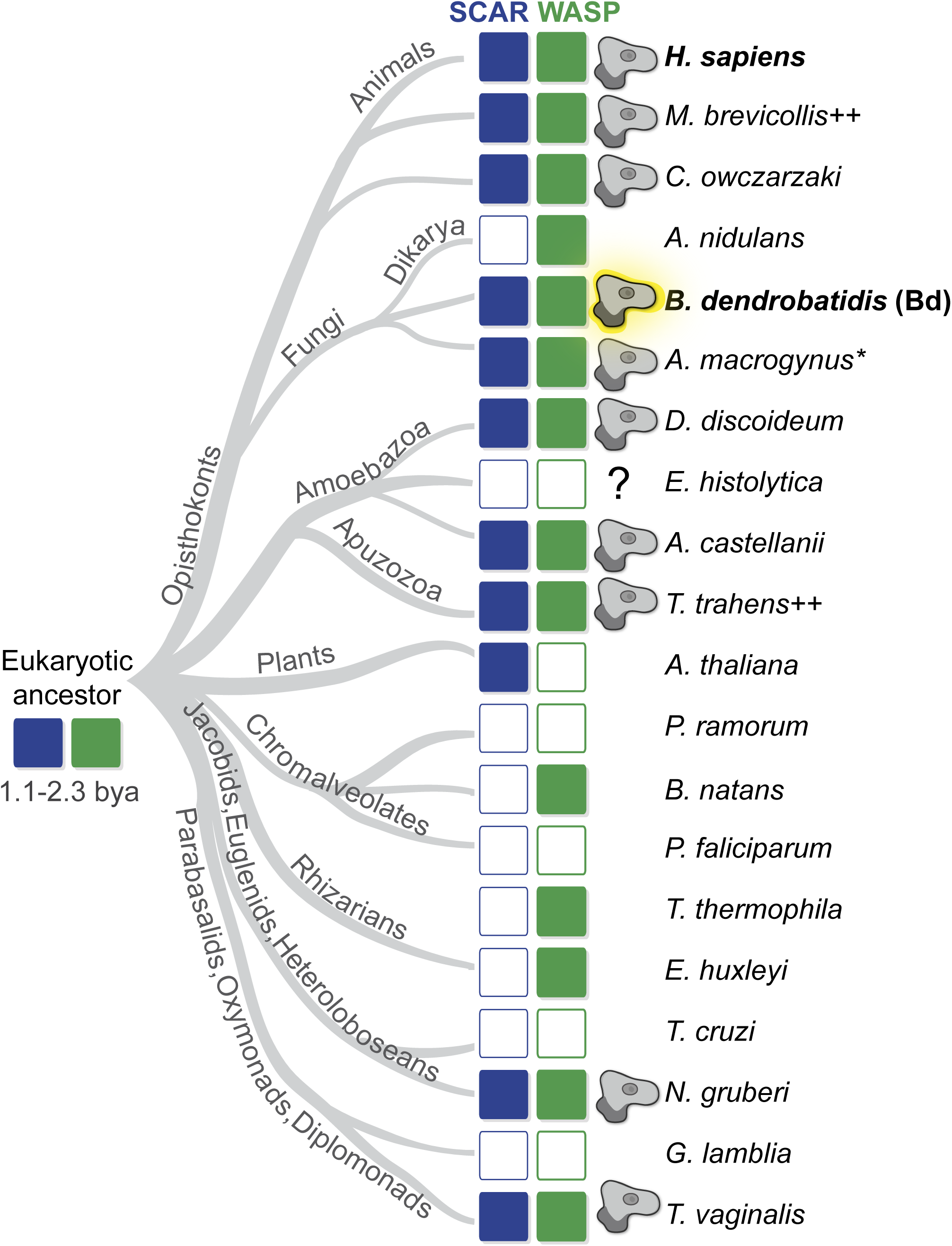
Only organisms that make pseudopods retain both WASP and SCAR genes. Diagram showing the relationships of extant eukaryotes (based on (Fritz-Laylin et al., 2010)), with presence or absence of SCAR (blue) and WASP (green) genes from complete genome sequences as described (Kollmar et al., 2012). The representative organism whose genome was used for the analysis is listed to the right. For groups with similar morphological and sequence patterns, a single species is used. For example, there is no known plant species that forms pseudopods or retains the WASP gene, so only a single species is shown (*Arabidopsis thaliana*); similarly, *Aspergillus nidulans* represents all dikarya. See (Kollmar et al., 2012) for additional sequence information. An amoeba glyph indicates organisms that build pseudopods. Outlined rectangles indicate a lack of identifiable gene. See Table 1 for citations and full species names. *Although we were not able to find a reference to pseudopod formation in *A. macrogynus*, a relative (*Catenaria anguillulae)* does assemble pseudopods used for motility (Gleason and Lilje, 2009; Deacon and Saxena, 1997). Because of this, and the conservation of both WASP and SCAR in *B. dendrobatidis* (highlighted in bold), we correctly predicted this species is also capable of pseudopod formation. ++These species form pseudopods for feeding, rather than motility. “?” Indicates uncertainty regarding the structure of the protrusions for phagocytosis in *Entamoeba histolytica* (see text). The time of divergence of extant eukaryotic groups has been estimated to be 1.1-2.3 billion years ago (Chernikova et al., 2011) (Parfrey et al., 2011; Knoll, 2014) and has been predicted to have possessed both WASP and SCAR gene families (Kollmar et al., 2012) and therefore may have built pseudopods.

Together with the metazoans and a handful of protists, fungi form a major clade known as the “opisthokonts” (Figure 8). Our identification of *α*-motility in a fungal species argues that the ancestor of all the opisthokonts was capable of fast, pseudopod-associated crawling and that multicellular fungi represent lineages that have lost *α*-motility (Fritz-Laylin et al., 2010). This loss appears to coincide with the disappearance from multicellular fungi of cell types lacking a cell wall. We propose that the limitations on pseudopod formation imposed by a rigid cell wall relieved the selection pressure to preserve gene networks specific to pseudopod formation, and the genes unique to this behavior were subsequently lost. The fungi would therefore represent a large eukaryotic lineage from which crawling motility has almost completely disappeared.

Images from earlier studies revealed individual Bd zoospores with irregular shapes and “cytoplasmic extensions” (Longcore et al., 1999). Actin-driven pseudopod formation and cell motility, however, was not previously described in Bd cells, in part because this species was discovered quite recently (Longcore et al., 1999) and relatively few studies have been devoted to its cell biology. In addition, Bd zoospores are quite small (<5 μm) and highly motile so that visualizing their tiny (∼1 μm) pseudopods requires physical confinement and high-resolution microscopy. Finally, because only recently released zoospores crawl, synchronization of cell cultures was crucial. These advances not only enabled us to observe *α*-motility, but also revealed Bd zoospores retracting their flagella by coiling the entire axoneme into the cell body in less than a second (**Video 5**), a process that has been observed to take minutes in other chytrid species (Koch, 1968).

The *α*-motility of Bd fills an important gap in our understanding of the life cycle of this pathogen. As proposed for other chytrid species (Gleason and Lilje, 2009), Bd zoospores may use pseudopods during the initial stages of their interaction with a host: either to move across epithelia or crawl between epithelial cells and invade the underlying stroma. Alternatively, our observation that newly hatched zoospores make more pseudopods suggests that Bd may rely on *α*-motility to crawl along or within the epithelial surface to uninfected tissues, or to exit the host.

Imaging chytrid zoospores provided key evidence for the involvement of WASP and SCAR in a conserved mode of cell migration, but further exploration of WASP and SCAR function in Bd is hampered by several factors. First, the zoospores are small enough to pose challenges to live-cell imaging. Second, the absence of genetic tools makes it impossible to fluorescently label or deplete proteins in live zoospores. Finally, the lack of potent and specific inhibitors of WASP and SCAR (Bompard et al., 2008; Guerriero and Weisz, 2007) precludes chemical disruption of their activity.

Several protein families are known to activate the Arp2/3 complex, including WASP, SCAR, WASH, JMY, and WHAMM (Rottner et al., 2010). A conventional explanation for the multiplicity of Arp2/3 activators is that each promotes construction of an actin network with a unique cellular function and/or location. However, the evolutionary connection between WASP, SCAR, and pseudopod formation suggests that nucleation promoting factors can work together, in this case to drive the explosive actin polymerization required for *α*-motility. Indeed, we find that WASP and SCAR colocalize at the leading edge of crawling neutrophils, and WASP depletion results in aberrant pseudopods and reduced motility, similar to reported effects of SCAR depletion (Weiner et al., 2006). But why have multiple distinct Arp2/3 activators rather than simply increasing the concentration of one of them? The answer may lie in the positive feedback that drives the explosive actin polymerization required for *α*-motility, but which can also result in spurious pseudopod formation (Chung et al., 2000). We propose that the use of two distinct activation systems—WASP and SCAR—reduces the probability of errant pseudopod formation by using both activators to raise Arp2/3 activity above the threshold required for robust pseudopod formation.

Under this “coincidence detection” model, an occasional spike in the local activity of either WASP or SCAR may be sufficient to trigger pseudopod formation, but activation of both should be more efficient at driving explosive actin assembly. This fits our observation that WASP depletion results in a subset of cells capable of pseudopod formation, but with less polymerized actin (Figure 3C–D) and reduced migration speeds (Figure 4B), and explains previous reports of incomplete penetrance of WASP and SCAR phenotypes (Veltman et al., 2012; Myers et al., 2005; Blagg et al., 2003; Weiner et al., 2006; Snapper et al., 2005). This model may also explain the rhino phenotype; without the additional Arp2/3 activation provided by WASP, the resulting sparse actin networks may collapse and coalesce into the observed horn-shaped structures. This idea is supported by studies of the upstream activators of WASP (Cdc42) and SCAR (Rac) that indicate that Rac mediates a positive feedback loop required for leading-edge formation, but the stability of the resulting protrusion requires Cdc42 (Srinivasan et al., 2003; Stradal and Scita, 2006).

This model also clears up confusion in the field regarding WASP’s role in cell migration. Our results are supported by papers showing that blood cells rely on WASP for efficient cell migration (Snapper et al., 2005; Kumar et al., 2012; Anderson et al., 2003; Blundell et al., 2008; Zhang et al., 2006; Zicha et al., 1998; Dovas et al., 2009; Worth et al., 2013; Binks et al., 1998), and others suggesting that WASP plays a direct role in protrusion formation, including pseudopods (Jones et al., 2002; Burns et al., 2001; Badolato et al., 1998; Jones et al., 2013; Shi et al., 2009; Ishihara et al., 2012). However, these data have been overshadowed by studies showing that fibroblasts do not require N-WASP for filopodia or sheet-like, surface-adhered lamellipodia (Snapper et al., 2001; Lommel et al., 2001; Sarmiento et al., 2008). Such papers have been cited as proof that all WASP family proteins are dispensable for protrusions in general (Small and Rottner, 2010). Such generalizations depend on two assumptions: that N-WASP and WASP have the same molecular function, and that the adherent motility of fibroblasts and *α*-motility use the same molecular pathways. However, recent molecular replacement studies have shown that WASP and the ubiquitously expressed N-WASP have different functions and cannot compensate for each other (Jain and Thanabalu, 2015). Furthermore, when one considers the large body of mammalian WASP literature in the light of distinct modes of motility, a simple pattern emerges: cell types that do not natively express WASP *do not make pseudopods* (although they may make surface-bound lamellipodia, linear filopodia, or adhesive structures called podosomes); WASP is *only* expressed in blood cells, and these cells use WASP for pseudopod-based migration. (See **Table S1** for an annotated summary of WASP/N-WASP literature.) The predominant view that WASP is not involved in cell migration demonstrates the peril of assuming that insights based on adhesion-dependent cell motility apply to other modes of cell crawling.

In addition to motility, Arp2/3 activators have been shown to play roles in other cellular processes, including endocytosis (Benesch et al., 2005; Merrifield et al., 2004; Naqvi et al., 1998). The relationship between cell motility and endocytosis and is complex and not completely understood (Schiefermeier et al., 2011; Traynor and Kay, 2007). Rapid pseudopod extension requires not only a large quantity of actin polymerization (Weiner et al., 2006), but also increases membrane tension (Diz-Muñoz et al., 2016), both of which counteract efficient clathrin- and actin-mediated endocytosis (Boulant et al., 2011). Despite this apparent dichotomy between protrusion formation and endocytosis, both SCAR and WASP protein families have been shown to interact with endocytosis pathways (Badour et al., 2007; Gautier et al., 2011). Accordingly, we find that WASP-deficient HL-60 cells maintain normal receptor internalization and recycling (Figure S3E), until pseudopod activity is activated by differentiation into neutrophils. After differentiation, in addition to being defective in building pseudopods, WASP-KD cells exhibit increased endocytosis and receptor recycling (Figure S3B-D). This is consistent with the idea that actin-mediated endocytosis is more efficient when cells are not making pseudopods.

Although a large number of eukaryotes make pseudopods (Table 1 and Figure 8), only two lineages are currently genetically tractable: animals and dictyostelids. Studies in both confirm that pseudopod formation involves both WASP (Jones et al., 2002; Burns et al., 2001; Badolato et al., 1998; Jones et al., 2013; Shi et al., 2009; Ishihara et al., 2012) and SCAR (Steffen et al., 2004; Miki et al., 1998; Veltman et al., 2012; Weiner et al., 2006), in contrast to some animal cell types that may only require SCAR for lamellipodia-based migration (Sarmiento et al., 2008; Bryce et al., 2005; Misra et al., 2007; Snapper et al., 2001). With our discovery of *α*-motility in the fungus *Bd*, we conclude that both proteins have been conserved together facilitate this evolutionarily ancient mode of cell motility.

## Materials and Methods

### Antibodies and Western Blotting

the rabbit anti-WASP antibody was from Santa Cruz (sc-8353), as was the goat anti-WAVE2 (sc-10394), the mouse anti-tubulin was from Sigma (DM1A), and the mouse anti-actin from Calbiochem (JLA20). Western blotting was conducted using standard protocols and horseradish-peroxidase-conjugated secondary antibodies (Jackson Laboratories).

### Generation of HL-60 cell lines

HL-60 lines were derived from ATCC #CCL-240, and were grown in medium RPMI 1640 supplemented with 15% FBS, 25 mM Hepes, and 2.0 g/L NaHCO3, and grown at 37C and 5% CO2. WASP-KD was achieved using Sigma’s Mission Control shRNA vector (TRCN0000029819, hairpin sequence: 5’-CCGGCGAGACCTCTAAACTTATCTACTCGAGTAGATAAGTTTAGAGGTCTCGTTTTT -3’), with corresponding control vector expressing anti-GFP shRNA (Catalog# SHC005, hairpin sequence: 5’-CCGGCGTGATCTTCACCGACAAGATCTCGAGATCTTGTCGGTGAAGATCTTTTT -3’). Lentivirus was produced in HEK293T grown in 6-well plates and transfected with equal amounts of the lentiviral backbone vector (either protein expression vector derived from pHRSIN-CSGW (Demaison et al., 2002), or shRNA expression vectors described above), pCMVΔ8.91 (encoding essential packaging genes) and pMD2.G (encoding VSV-G gene to pseudotype virus). pHRSIN-CSGW and packaging vectors were obtained from the laboratory of Ron Vale. After 48hr, the supernatant from each well was removed, centrifuged at 14,000 g for 5 min to remove debris and then incubated with ∼1×10^6^ HL-60 cells suspended in 1 mL complete RPMI for 5-12 hours. Fresh medium was then added and the cells were recovered for 3 days to allow for target protein or shRNA expression. TagRFPt-WASP fusion was cloned by first swapping out eGFP for TagRFP-T (Shaner et al., 2008) in the pHRSIN-CSGW by PCR amplifying TagRFP-T with 5’CCCGGGATCCACCGGTCGCCACCATGGTGTCTAAGGGCGAAGAGCTGATTAAG G3’ and 5’GAGTCGCGGCCGCTTTAACTAGTCCCGCTGCCCTTGTACAGCTCGTCCATGCCATTAAGTTTGTGCCCC3’ primers, and cloning the resulting PCR product into pHRSIN-CSGW using NotI and BamHI to produce the pHR-TagRFP-T vector. Then, the WASP open reading frame was PCR amplified from cDNA (NCBI accession BC012738) using 5’GCACTAGTATGAGTGGGGGCCCAATGGGAGGAA3’ and 5’AAGCGGCCGCTCAGTCATCCCATTCATCATCTTCATCTTCA3’ primers, and cloned into the pHR-TAGRFP-T backbone using NotI and SpeI to result in a single open reading frame containing TagRFPt, a flexible linker (amino acids GSGTS), followed by full length WASP. The WASP shRNA rescue vector was cloned by inserting a P2A cleavage site (Kim et al., 2011) between the linker and a WASP open reading frame edited with site-directed mutagenesis to contain three silent mutations within the shRNA-targetting region (5’cgagacctctaaacttatcta3’ was changed to 5’CGAaACCTCTAAgCTcATCTA3’, with silent mutations in lower case). A corresponding control vector was designed to express TagRFPt with the flexible linker, but no portion of WASP. The Hem1-YFP line (R6) was previously described (Weiner et al., 2007), and was developed by selecting HL-60 cells expressing both an shRNA targeting native Hem1 and a YFP-tagged version of Hem-1, allowing fluorescence imaging of the SCAR complex in HL-60 cells. shRNA lines were selected by puromycin (1 μg/mL for at least 1 week), and florescent cell lines by FACS. HL-60 cells were differentiated by treatment with 1.3% DMSO for 5 days.

### Cytometry

Fluorescence-activated cell sorting (FACS) analysis was performed on a FACSCalibur analyzer (Bd), Data were analyzed with FlowJo software (Tree Star) and dead cells were gated out using forward and side scatter for all analyses. A FACS Aria II was used for sorting. All FACS analysis was performed at the Laboratory for Cell Analysis (UCSF).

### Imaging

EZ-TAXIScan (Effector Cell Institute, Tokyo) analysis of HL-60 cell migration between glass surfaces was conducted as previously described (Millius and Weiner, 2010), and cell migration analyzed using Chemotaxis and Migration Tool (Ibidi). Fixed HL-60 cells were imaged with a 100× 1.40 NA oil PlanApo objective on a motorized inverted microscope (Nikon Ti-E) equipped with a spinning disk (Yokogawa CSU22) and EMCCD camera (Photometrics Evolve). Live TIRF images were acquired by plating HL-60 cells on coverglass cleaned by a 30 min incubation in 3M NaOH, followed by four washes with PBS, pH 7.2, coated for 30 min with 100 μg/mL bovine fibronectin (Sigma F4759) resuspended in PBS. TIRF microscopy images were acquired on a Nikon TE2000 inverted microscope equipped with a 1.45 NA oil 60× or 100× PlanApo TIRF objective and an EMCCD (Andor iXon+), using previously described imaging conditions (Weiner et al., 2007), briefly, differentiated HL-60 cells were plated on fibronectin-coated coverslips in modified HBSS supplemented with 0.2% serum albumin and imaged with 100 ms exposures every 1–2 s. Fixed chytrid cells were imaged using an inverted microscope (Nikon Ti-E, Tokyo, Japan) equipped with a spinning-disk confocal system with 33 μm pinholes and a 1.8× tube lens (Spectral Diskovery), a Nikon 60× 1.45 NA Apo TIRF objective, and a CMOS camera (Andor Zyla 4.2). DIC microscopy was performed on an inverted microscope (Nikon Ti-E) with a light-emitting diode illuminator (Sutter TLED) and a Nikon 100× 1.45 NA Apo TIRF objective; images were acquired on a CMOS camera (Andor Zyla 4.2). All microscopy hardware was controlled with Micro-Manager software (Edelstein et al., 2010). Image analysis was performed with the ImageJ bundle Fiji (Schindelin et al., 2012). All imaging was done at room temperature.

### Quantitation of actin polymerization by flow cytometry

HL-60 cells were depolarized in serum free medium supplemented with 2% low-endotoxin BSA (Sigma) for 1 hour at 37C and 5% CO2 before simulation with 20nM fMLP for the indicated time. Cells were immediately fixed with 4% paraformaldehyde in cytoskeleton buffer on ice for 20 min, stained with PBS supplemented with 2% BSA, 0.1% Triton X-100, and 66 nM Alexaflor-488 conjugated phalloidin (Molecular Probes, catalog # A12379) for 20 min, and washed thrice with PBST before FACS analysis.

### Cell adhesion assay

Differentiated control and WASP-KD HL-60 cells were each stained with either green or blue acetoxymethyl (AM) ester dyes (CellTrace calcein green and blue, ThermoFisher) and equal numbers mixed and allowed to attach to fibronectin-coated coverglass-bottomed 96 well plates for 30 min at 37C. One set of wells was gently washed three times with fresh media. 100 random locations within the well were immediately imaged, and the percent remaining cells calculated and normalized to control unwashed wells.

### Transferrin uptake endocytosis assays

5×10^6^ differentiated HL-60 cells were washed twice with ice-cold serum-free growth medium (SF), transferred to 37C for 5 min (to clear surface-bound transferrin), and chilled on ice for 1 min. An equal volume of cold SF supplemented with 100ug/mL Alexa488 conjugated transferrin (McGraw and Subtil, 2001) (Molecular Probes T-13342) was added, and incubated on ice for 10 min. Cells were then washed twice with cold SF medium, transferred to 37C for the indicated time period, washed twice with ice-cold acid buffer (8.76 g NaCl, 9.74 g MES in 900 mL, pH to 4.0, water to 1 L), fixed in 4% paraformaldehyde in 1x PBS for 20 min, and washed twice more with ice-cold PBS before immediate FACS analysis.

### Motility of Chytrid zoospores

*Batrachochytrium dendrobatidis* strain JEL423 was obtained from the laboratory of Joyce Longcore (University of Maine), and grown in 1% tryptone broth or on agar plates (1% tryptone, 2% agar) at 25C. Before imaging, cultures were synchronized by either three washes in 1% tryptone, and zoospores harvested 2 hours later (for liquid cultures), or flooding agar plates with ∼2 mL water (for agar plates), passed through a 40 μm filter (Falcon), and collected by centrifuging at 1200 g for 5 min, and resuspended in Bonner’s Salts (Bonner, 1947). Cell motility was imaged by sandwiching cells between a #1.5 glass coverslip and glass slide (cleaned by sonicating in pure water) separated using 1 μm glass microspheres (Bangs Laboratories). Coverslip and glass slide were sonicated in deionized water, and dried. Cells were treated with either 10 μM CK-666 (Sigma) or 10 nM latrunculin B (Sigma) while cells were adhered to Concalavin-A coated glass. For visualization of polymerized actin: 400 μL fixation buffer (50 mM cacodylate buffer, pH 7.2) supplemented with 4% gluteraldehyde, was added to 100 μL cells attached to a Concalavin-A coated coverslip, and incubated for 20 min at 4C. Samples were quenched with tetraborohydride, permeabilized with 0.1% Triton X-100, incubated for 20 min with Alexa488-labeled phalloidin (Invitrogen), rinsed 4 times, and imaged as above. Samples for scanning electron microscopy were fixed as above, stained with osmium tetroxide, dehydrated, critical point dried, and Au/Pd sputter coated according to standard protocols and imaged using a Hitachi S-5000 scanning electron microscope in the University of California, Berkeley Electron Microscopy Laboratory.

### Summary of supplemental material

The supplemental material includes 4 supplemental figures: S1: WASP localizes to pseudopods of migrating neutrophils, S2: WASP depletion in neutrophils leads to dynamic rhino protrusions, S3: WASP is not required for adhesion or endocytosis by HL-60 cells, and S4: additional examples of chytrid pseudopods. It also includes 5 videos: 1: TIRF microscopy of HL-60 cells, 2: chemotaxis of HL-60 cells, 3: time lapse movies of chytrid zoospores making pseudopods, 4: chrytrid cells crawling, and 5: an example of a chytrid zoospore retracting its flagellum. An annotated table summarizes the literature on the roles of WASP and N-WASP in protrusion formation and cell motility.

## Acknowledgements

We would like to thank Jasmine Pare for help with cell culture and HL-60 cell migration assays, and Joyce Longcore, Jason Stajich, and Erica Rosenblum for strains and helpful discussions. This work was supported by the National Institute of General Medical Sciences of the National Institutes of Health (R01-GM061010 to R.D.M.), by the Howard Hughes Medical Institute Investigator program (R.D.M.), and by a postdoctoral fellowship from the Helen Hay Whitney Foundation supported by the Howard Hughes Medical Institute (L.K.F.-L.). The authors declare no competing financial interests.

### Author Contributions

Lillian K. Fritz-Laylin: conceptualization, project administration, funding acquisition, methodology, investigation, formal analysis, validation, visualization, writing (original draft preparation), and writing (review & editing). Samuel J. Lord: methodology, investigation, formal analysis, validation, visualization, writing (original draft preparation), and writing (review & editing). R. Dyche Mullins: funding acquisition, supervision, and writing (original draft preparation, review & editing).

---

**Figure S1.**
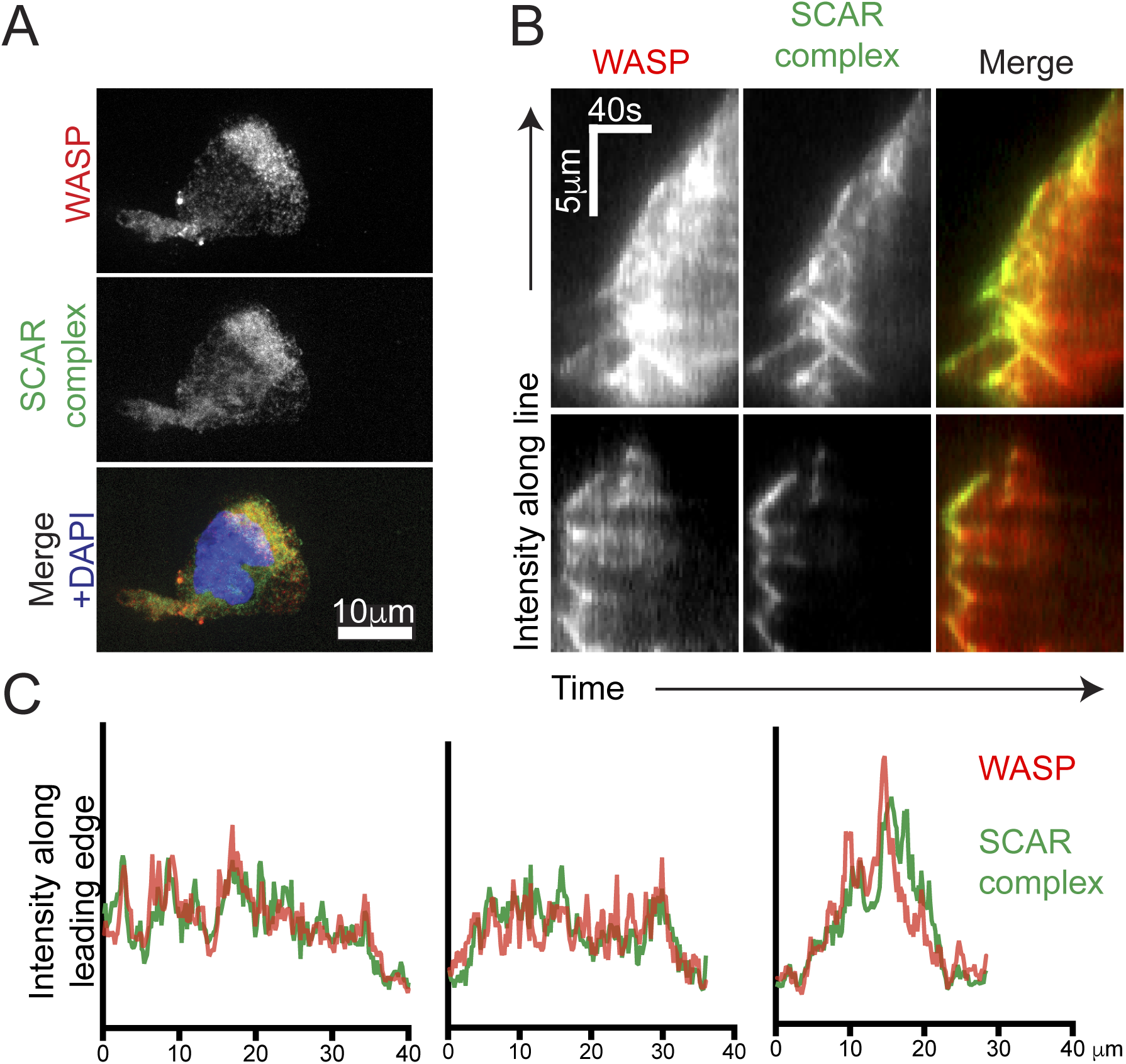
WASP localizes to pseudopods of migrating neutrophils. **(A)** Maximum projection image of localization of TagRFP-WASP and Hem-1-YFP (a member of the SCAR regulatory complex) expressed in differentiated HL-60 cells fixed while migrating on a fibronectin-coated surface. **(B)** Kymographs of pseudopods of two live HL-60 cells, showing colocalization over time of TagRFP-WASP and Hem-1-YFP patterns. Time from left (T=0) to right, and direction from bottom (inside cell) to top (outside cell). See **Video 1** for the cells from which these kymographs were generated. **(C)** Linescans following the contour of the leading edge of each individual cell shown Figure 1A-B, showing that TagRFP-WASP (red) partially colocalizes with Hem-1-YFP (green).

**Figure S2.**
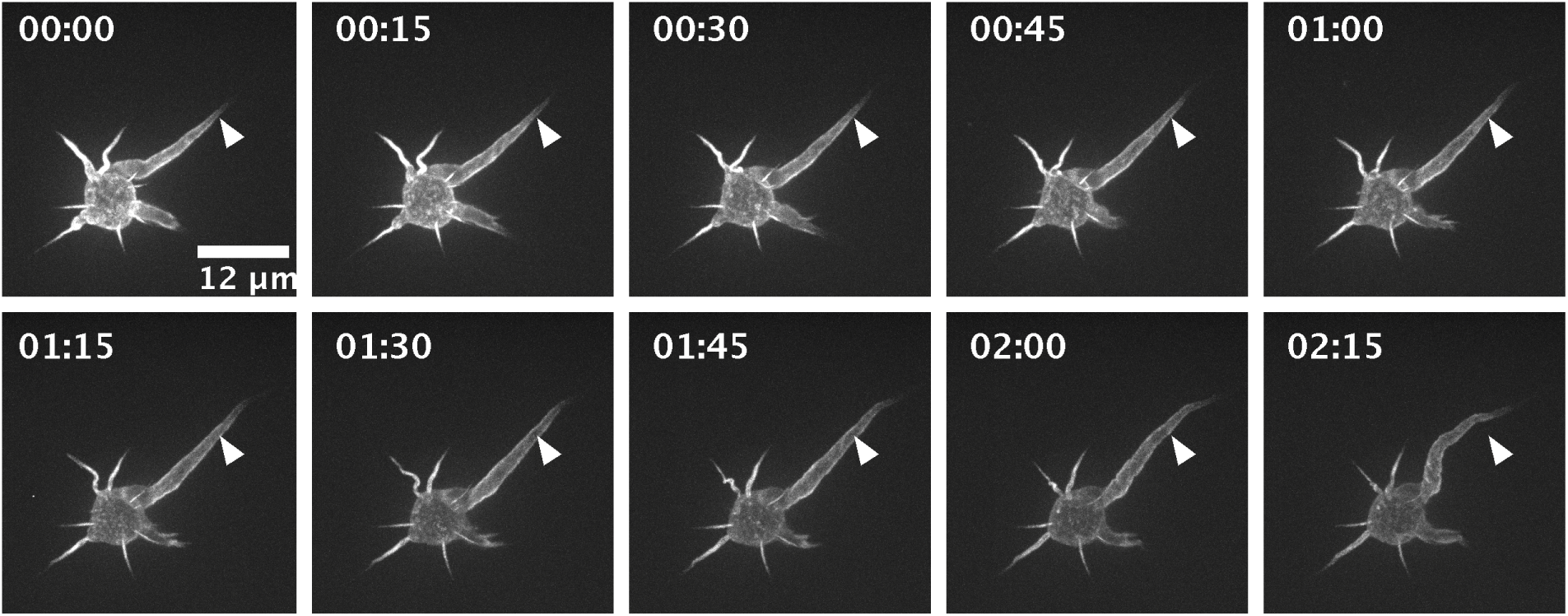
WASP depletion in neutrophils leads to dynamic rhino protrusions. Time-lapse microscopy of a live WASP-KD HL-60 cell with aberrant “rhino” protrusions, with polymerized actin visualized using Utrophin261-mCherry. Time is shown in (min: seconds), and an immobile arrowhead highlights protrusion dynamics.

**Figure S3.**
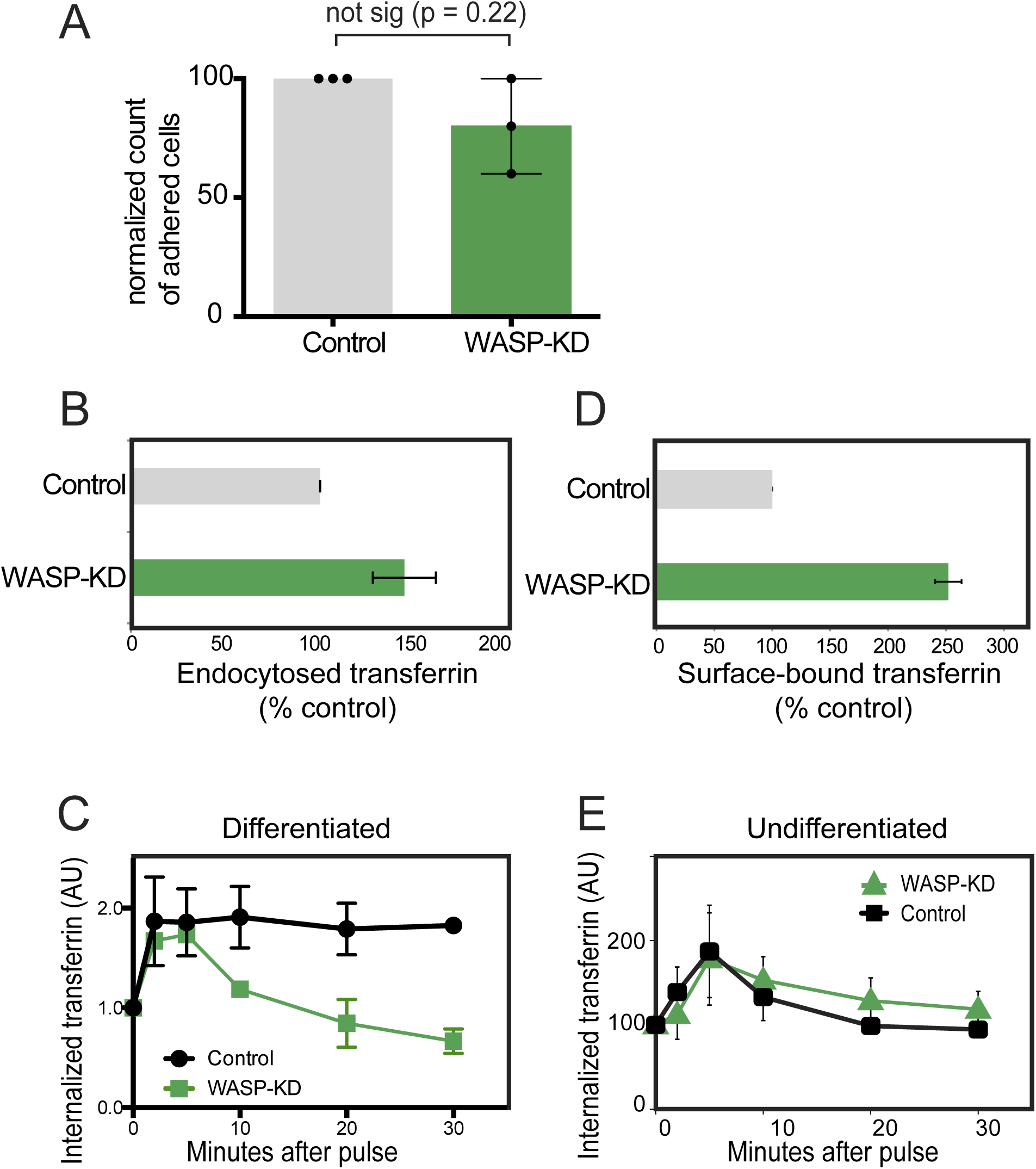
WASP is not required for adhesion or endocytosis by HL-60 cells. **(A)** WASP-KD does not significantly reduce the HL-60 cells’ ability to adhere to fibronectin-coated surfaces. Bars represent averages from three biological replicates, normalized to an internal control in each experiment; p value from two-tailed paired t-test. At least 1000 cells were counted for each internal control. **(B)** Steady state endocytosis was measured for WASP-KD (green) and control cells (black) by incubating cells for 10 minutes at 37C with fluorescent transferrin, immediately washing with ice-cold acid buffer (to remove surface-bound transferrin). Cells were then fixed, and endocytosed tranferrin quantified by FACS analysis. Values were normalized to percent control within each of three independent experiments, with 10,000 cells analyzed for each sample‥ **(C)** Quantitation of actin-mediated endocytosis and receptor recycling in differentiated (neutrophil-like) control (black circles) and WASP-KD HL-60 cells (green squares). Cells were incubated with florescent transferrin, placed at 37C for the indicated time, washed with ice-cold acid buffer (to remove surface-bound transferrin), and fixed for FACS analysis. Within each experiment, samples for each cell line were normalized to time zero. Averages and standard deviations for three independent experiments are shown, with 10,000 cells analyzed for each sample. (**D**) Transferrin receptor density was measured by incubating cells at 37C in serum-free medium (to remove surface-bound transferrin), chilling cells and incubating on ice with florescent transferrin. Cells were then washed with PBS to remove unbound transferrin and fixed. Surface-bound transferrin was then quantitated by FACS analysis. Values were normalized to percent control within each of three independent experiments, with 10,000 cells analyzed for each sample. (**E**) Quantitation of actin-mediated endocytosis and receptor recycling in undifferentiated control (black squares) and WASP-KD HL-60 cells (green triangles). Cells were incubated with florescent transferrin, placed at 37C for the indicated time, washed with ice-cold acid buffer (to remove surface-bound transferrin), and fixed for FACS analysis. Within each experiment, samples for each cell line were normalized to time zero. Averages and standard deviations for three independent experiments are shown, with 10,000 cells analyzed for each sample. Note that the recycling assays are normalized. For comparing absolute amount of material endocytosed, refer to (B).

**Figure S4.**
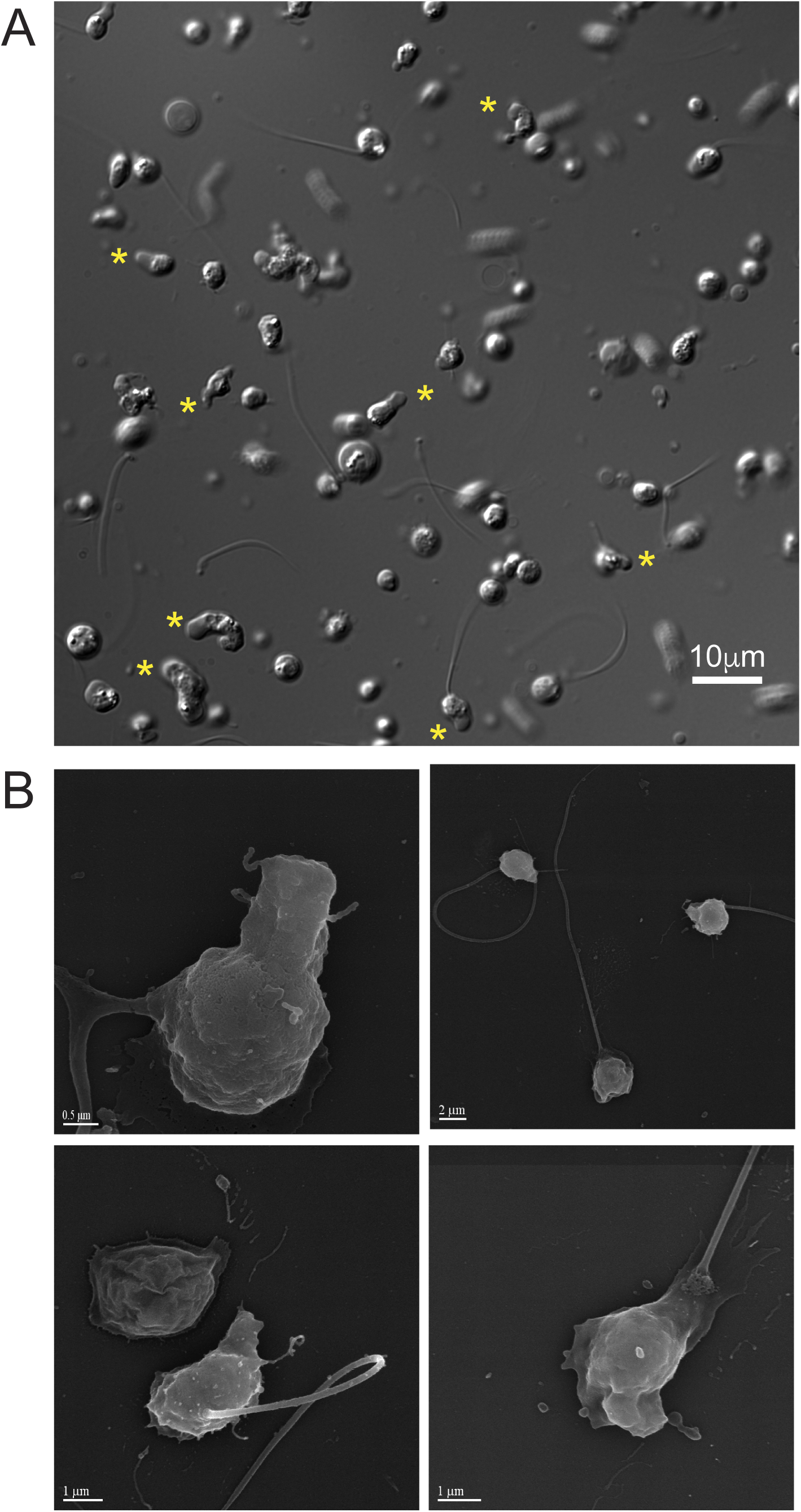
Additional examples of chytrid pseudopods. **(A)** DIC image of representative field of synchronized Bd zoospores. Asterisks are highlighted by cells with obvious pseudopods. Rapidly-swimming flagellate cells are blurred. **(B)** Additional examples of scanning electron micrographs of chytrid zoospores.

## Video Legends

**Video 1.** Five examples of lapse movies showing TIRF microscopy of live differentiated HL-60 neutophil cells expressing Hem-1-YFP (top, and green in overlay) and TagRFP-WASP (middle, and red in overlay). White lines indicate position for kymographs in Figure S1B.

**Video 2.** Migration of control differentiated HL-60 neutrophil cells and of differentiated WASP-KD HL-60 cells in 2D environment (EZ-TAXIScan assay). Cells expressing control shRNA migrating in a chemoattractant gradient (source is at top) between two glass surfaces with 5 μm spacing. All cells initially within the field of view were manually tracked, as shown. Images were acquired every 20 seconds and are displayed at 300X speed. Many WASP-KD cells exhibited the rhino phenotype and their motility was effectively abolished (e.g. cells 11, 32, 33, 39 and 42).

**Video 3**. Time lapse movies showing the representative examples of Bd chytrid zoospores with pseudopods imaged using DIC microscopy pictured in Figure 4B, including a cell with flagella (bottom left), a cell without a flagellum (bottom right), and one cell of each (top).

**Video 4**. Examples of Bd chytrid cells crawling between two glass coverslips separated by 1 μm glass beads. See also Figure 5.

**Video 5**. Time lapse imaging showing an example of flagellar retraction by Bd chytrid zoospores.

